# Skeletons in the closet? – Using a bibliometric lens to visualise phytochemical and pharmacological activities linked to *Sceletium*, a mood enhancer

**DOI:** 10.1101/2023.08.11.552916

**Authors:** Kaylan Reddy, Gary I. Stafford, Nokwanda P. Makunga

**Affiliations:** Department of Botany and Zoology, Private Bag X1, Natural Sciences Faculty, Matieland, Stellenbosch University South Africa; Department of Plant and Soil Sciences, University of Pretoria, Pretoria, South Africa

**Keywords:** alkaloid chemistry, central nervous system activity, kanna, secondary metabolites, pharmacology, phytochemistry

## Abstract

Plants from the *Sceletium* genus (Aizoaceae) have been traditionally used by the Khoe-Sān people in southern Africa, mainly for thirst and hunger relief, pain reduction and spiritual purposes, particularly *Sceletium tortuosum*. The research on this species has seen rapid growth with advancements in analytical and pharmacological tools. The Web of Science (WoS) database was searched for articles related to ‘Sceletium’ and ‘Mesembrine’. These data were additionally analysed by bibliometric software (VOSviewer) to generate term maps and author associations. The thematic areas with the most citations were, South African Traditional Medicine for mental health (110) and anxiolytic agents (75). Pioneer studies in the genus focused on chemical structural isolation, purification and characterization and techniques such as thin layer chromatography, liquid chromatography (HPLC, UPLC and more recently, LC-MS), gas chromatography mass spectrometry (GC-MS) and nuclear magnetic resonance (NMR) to study mesembrine alkaloids. Different laboratories have used a diverse range of extraction and pre-analytical methods that become routinely favoured in the analysis of the main metabolites (mesembrine, mesembranol, mesembranone and Sceletium A4) in their respective experimental settings. In contrast with previous reviews, this paper identified gaps in the research field, being a lack of toxicology assays, a deficit of clinical assessments, too few bioavailability studies and little to no investigation into the minor alkaloid groups found in *Sceletium*. Future studies are likely to see innovations in analytical techniques like leaf spray mass spectrometry and direct analysis in real-time ionization coupled with high-resolution time-of-flight mass spectrometry (DART-HR-TOF-MS) for rapid alkaloid identification and quality control purposes. While *S. tortuosum* has been the primary focus, studying other *Sceletium* species may aid in establishing chemotaxonomic relationships and addressing challenges with species misidentification. This research can benefit the nutraceutical industry and conservation efforts for the entire genus. At present, little to no pharmacological information is available in terms of the molecular physiological effects of mesembrine alkaloids in medical clinical settings. Research in these fields is expected to increase due to the growing interest in *S. tortuosum* as a herbal supplement and the potential development of mesembrine alkaloids into pharmaceutical drugs.

## Introduction

The plant *Sceletium tortuosum* (L.) N.E.Br. has well-documented medicinal activity and ethnopharmacology (Smith et al., 1998; Gericke and Viljoen, 2008). The most popular and well-known taxa from the *Sceletium* genus (Family: Aizoaceae, subfamily: Mesembryanthemoideae) is *Sceletium tortuosum*. *S. tortuosum* is also referred to as kanna, channa, kougoed or ‘sceletium’ (Smith et al., 1998). This species is a climbing or creeping perennial with succulent leaves and stems that become thick and slightly woody with age (Klak et al., 2007). Species are distinguished based on morphology. An important diagnostic feature of this genus is the skeletonised veins that are apparent when leaves dry (Fig. 1a). The typical growth form exhibits a scandent nature (Fig. 1b and c) together with leaves that have idioblasts or “bladder cells” (Fig. 1d). The flower colour of petals ranges from white, yellow to pale pink (Fig. 1e). The seeds of *Sceletium* species are brown to black kidney-shaped and these are small in diameter ranging from 1-2 mm. (Fig. 1f).

**Fig. 1.**
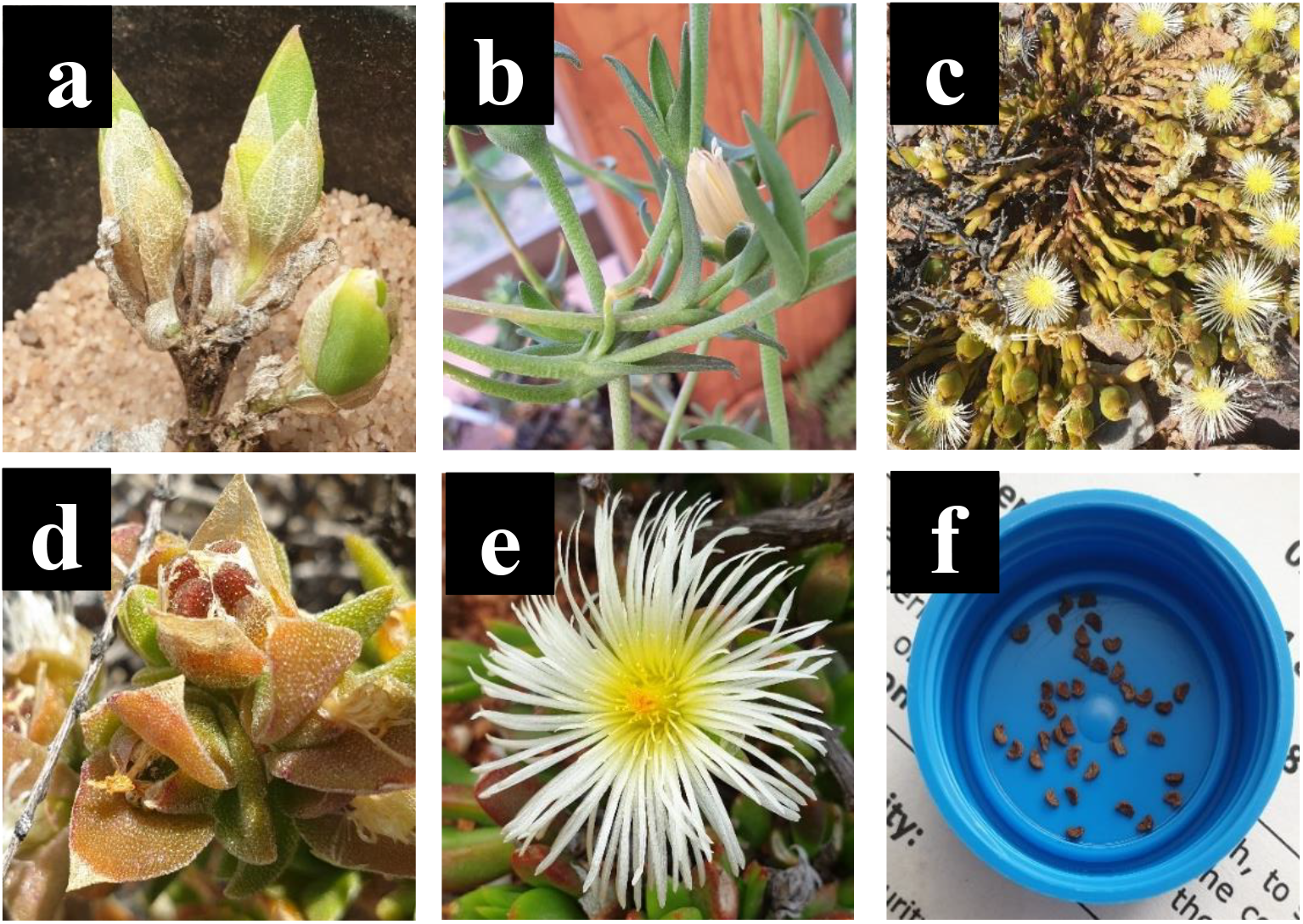
(a) *Sceletium rigidum*; (b) Image of *Sceletium subvelutium* (Syn. *Mesembryanthem varians*); (c) climbing or decumbent habit form of growth; (d) characteristic idioblasts (bladder-like cells) on *Sceletium* leaves; (e) flower structure of *Sceletium* species and (f) characteristic kidney-shaped seeds. (All images taken by N Makunga and K Reddy)

The plant is indigenous to southern Africa where it has been traditionally used in folk medicine by the Khoekhoen and Sān (Khoe-Sān/KhoiSan) people as a masticatory agent or as a mood elevator (Gericke and Viljoen, 2008). More recently, *S. tortuosum* has been commercialised as an anti-depressant or anxiolytic and it is also recommended for attention deficit disorders, as it aids in mental alertness (Harvey et al., 2011). The chemical constituents which were recognized for their medicinal activity are a group of mesembrine alkaloids that are uniquely associated with *Sceletium* taxa, however, they do share some similarities with Amaryllidaceae alkaloids. There has been a particular emphasis on mesembrine (Fig. 2a), mesembrenone (Fig. 2b) and Δ^7^mesembrenone (Fig. 2c) as biomarker compounds due to more scientific information being available in terms of chemical characterization. As a result, many producers of various phytopharmaceuticals, derived from *S. tortuosum* in particular, utilise these compounds in their quality assurance profiling regimes as part of the production value chain linked to this species. The industry based on this particular species is ever growing as more scientific information validating the central nervous system (CNS) effects of the plant populates the primary literature. Thus far, there have been several comprehensive reviews based on the chemistry of alkaloids found in *Sceletium* (Jeffs et al., 1982; Lewis, 1995, 2001; Jin, 2016; Jin and Yao, 2019). Although this list may not necessarily be comprehensive as it is based on a Scopus database search, other reviews that focus on *Sceletium* and its phytochemistry and pharmacology include the work of Gericke and Viljoen, (2008); Stafford et al. (2008); Van Wyk, (2011, 2015); Krstenansky, (2017); Makolo et al. (2019); Faro et al. (2020). These reviews discuss 1) the ethnobotanical history and chemical diversity in the genus (Smith et al., 1998); 2) the pharmacological and chemical evidence of ethnobotanical use in *Sceletium* (Gericke and Viljoen, 2008); 3) plants from South Africa with CNS-effects used for mental health purposes (Stafford et al., 2008); 4) the commercial potential of medicinal plants in South Africa (Van Wyk, 2011, 2015); 5) the occurrence, chemistry and pharmacology of mesembrine alkaloids (Krstenansky, 2017); 6) the distribution, structural elucidation, biosynthesis, organic synthesis, chemotaxonomy and biological activities of (-)-mesembrine from *Sceletium* species (Makolo et al., 2019); and, 7) the biomedical activities of new psychoactive substances from natural origins (Faro et al., 2020). Within this current paper, we provide an update on analytical techniques used to study *Sceletium tortuosum* and its relatives, where possible. We also summarise studies that focus on chemical variation as much quantitative and qualitative information is still presently missing with regards to the biochemical components that make up the phytochemical profiles of these plants. This paper also presents findings on the use of VOSviewer to identify gaps and trends in *Sceletium* research, which may be of value for other scientists and industry to decide on areas to research within the available options. Furthermore, there is great interest in the use of *Sceletium* species and *Sceletium* alkaloids against anxiety (Shikanga et al., 2011; Loria et al., 2014a) and depression (Gericke and Viljoen, 2008; Krstenansky, 2017) but preclinical and clinical evidence that validate these particular applications, that are grounded in an ethnobotanical context, are still limited. In spite of this, this has led to the commercialisation of *S. tortuosum* for various phyto-pharmaceutic markets (Patnala and Kanfer, 2013; Krstenansky, 2017).

**Fig. 2.**
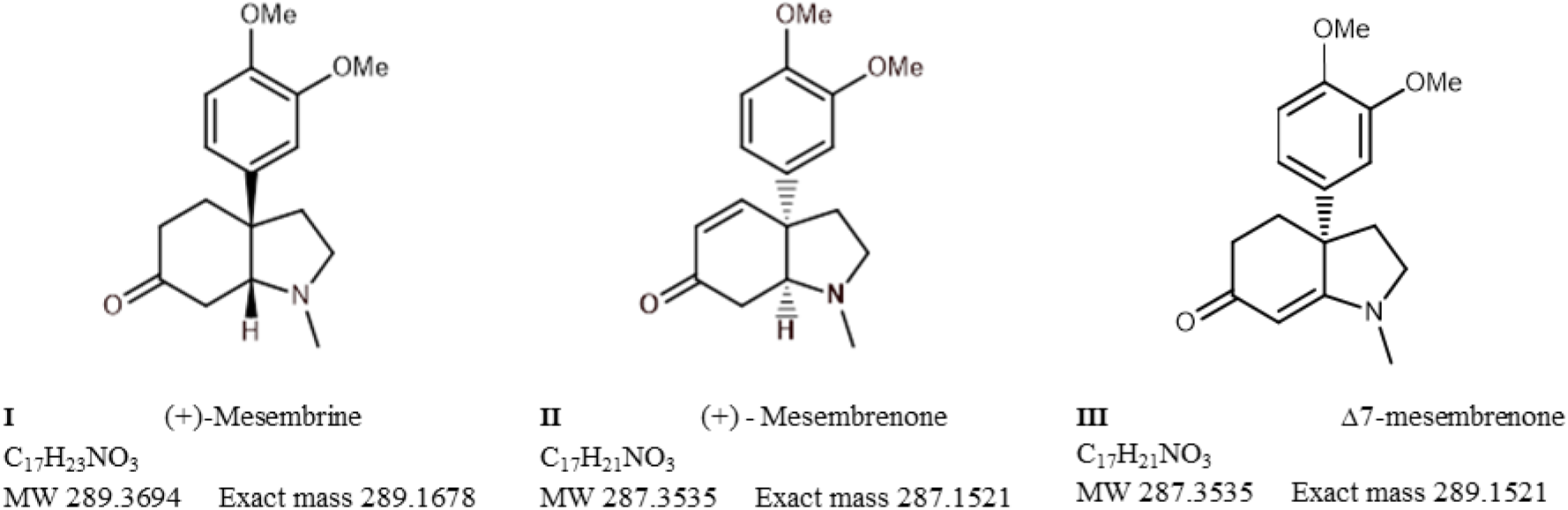
Chemical structures of mesembrine, mesembrenone and Δ^7^mesembrenone of greatest medicinal importance in *Sceletium* research

In order to get an overview of the available literature, a systematic bibliometric analysis was undertaken. Currently, there is a great deal of chemical and pharmaceutical studies on *Sceletium* but aspects linked to the taxonomy and geographical occurrence of many species of *Sceletium* apart from *S. tortuosum* are limited. This is of relevance as species misidentifications and biodiversity losses may prevail. The first part of this review thus aimed to collate information linked to the taxonomy and distribution of *Sceletium* species. These data were collected from databases such as SANBI-BODATSA and iNaturalist as an introduction before an update on the pharmacology and chemistry observed within the genus is presented. It is imperative to prioritize the correct collection of species and as such an understanding of the taxonomy of the genus should be consulted. The current trends within the literature and associated authors on a global scale. The present review summarises the studies conducted on the *Sceletium* genus and its chemical constituents over time in terms of the progress in phytochemistry, ethnobotanical use and pharmacology. This work intends to expose the current gaps within *Sceletium* research. Here, we report on studies from 1961 to the present and direct attention to recent advancements and future directions that may further develop quality, safety, and toxicological standards for therapeutic and nutraceutical applications concerning *S. tortuosum* and its relatives.

### Taxonomy and Distribution

The species currently recognised are *S. crassicaule* (Haw.) L. Bolus, *S. emarcidum* (Thunb.) L. Bolus ex H.J. Jacobson, *S. exalatum* Gerbaulet*, S. expansum* (L.) L. Bolus, *S. rigidum* (Fig. 1A), L. Bolus, *S. strictum* L. Bolus, *S. tortuosum* and *S. varians* (Haw.) Gerbaulet (Fig. 1B), as revised by Gerbaulet (Gerbaulet, 1996). Several species were reduced to being combined into the same species including *S. joubertii* L. Bol. And *S. namaquense* L. Bol., now considered to be part of the *S. tortuosum* complex. Taxonomically the plant genus was established in 1925 by N.E. Brown. However, it should be noted that Klak et al. (2007), in their phylogenetic study of the family, proposed that Mesembryanthemoideae should consist of the single genus *Mesembryanthemum*. Thus *Sceletium* was reduced to synonymy to *Mesembryanthemum*, thus the eight species of *Sceletium* (above) are currently accepted as *Mesembryanthemum crassicaule* Haw.*, M. emarcidum* Thunb.*, M. exalatum* (Gerbaulet) Klak*, M. expansum* L.*, M. archeri* (L.Bolus) Klak (*=S. rigidum*)*, M. ladismithiense* Klak (*=S. strictum*)*, M. tortuosum* L. and *M. varians* Haw. However, for the purpose of this particular review, *Sceletium* is used as this is still predominantly used in industry, in scientific works on the commercially important *Sceletium tortuosum*, particularly related to its chemistry and pharmacology, and non-scientific settings.

As part of this review, a distribution of *Sceletium* species was generated from the SANBI-BODATSA (South African National Biodiversity Institute - Botanical Database of Southern Africa), this database contained information sourced from observational data, herbaria, literature, collector information and species checklists. The majority of the observations were in the Western Cape of South Africa with some in the Northern and Eastern Cape provinces as illustrated in Fig. 3. A particular emphasis has been placed on *S. tortuosum* in the literature for its medicinal properties. The distribution of *S. tortuosum* has been reported in the southwestern areas of South Africa (Gericke and Viljoen, 2008). The plant has an affinity for arid environments and has been reported to grow from Namaqualand through to Aberdeen in South Africa (Chesselet, 2005).

**Fig. 3.**
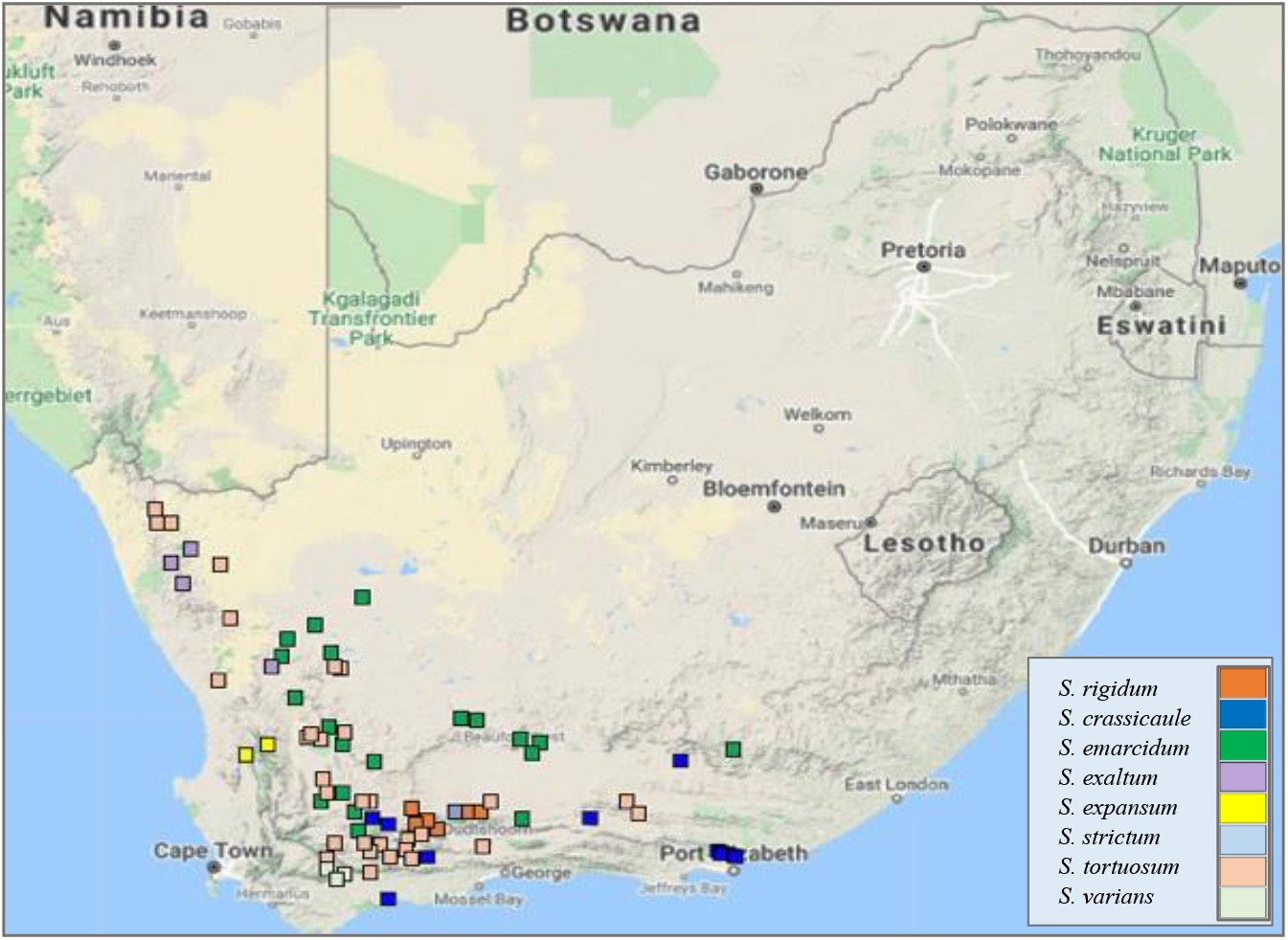
A: Geographic distribution of wild collections of eight species of *Sceletium* in South Africa (Data obtained from SANBI-BODATSA Database)

### Ethnobotany

Simon van der Stel’s, the last commander and first Governor of the Dutch Cape Colony, journey to Coperbergh (near present-day Okiep and Carolusberg, in the Northern Cape, South Africa) in 1685 made note of how *kanna* was consumed by the native people and details of its processing were included in the descriptions related to the species. The journal had the following quotation (translated from Dutch):

“They chew mostly a certain plant which they call Canna and which they bruise, roots as well as the stem, between the stones and store and preserve in sewn-up sheepskins”.

Between the date ranges of 1772 and 1774, a Swiss botanist and student of Linnaeus, Carl Peter Thunberg, made journeys to the Eastern Cape and reported on the value of the sedative plants that were found in the locality of present-day Oudtshoorn in the Little Karoo, South Africa (Gordon, 1996). At this time, it was reported that the land was inhabited by the Attaqua Khoikhoi (Khoekhoen), who called the land ‘Cannaland’ (Gordon, 1996). Several authors reported on the use of *Sceletium tortuosum* and *Sceletium expansum* by the indigenous people of South Africa. Some reported that the plants were used as tinctures (Pappe, 1868), occasionally snuffed or smoked, as teas (Jacobson, 1960; Smith et al., 1996; Van Wyk and Wink, 2018), or just purely as a form of recreation more than a medicine (Hartwich and Zwicky, 1914). Watt and Breyer-Brandwijk (Watt and Breyer-Brandwijk, 1962) indicated that in Namaqualand both the aerial and the underground (root) parts were used to make *kougoed* and how *Sceletium tortuosum* was used as an agent to help with pain, hunger relief, cholic and restlessness in infants by the Nama people (Watt and Breyer-Brandwijk, 1962). Since the review paper of Smith et al. (1998), an increasing body of scientific information, associated in particular with *Sceletium tortuosum*, has emerged. However, scientific interest in this species is leading to continuous progress in the areas of phytochemistry and pharmacology. This review aimed to provide visual networks linked to past research and identified current trends. We provide a historical account of the use of analytical techniques and pharmacological bioassays that have been employed to study *S. tortuosum* and its relatives. Finally, gaps in knowledge, recommendation and best practice in studying these neurologically acting medicinal plants is presented.

### Bibliometric analysis

#### Data Sources

The Web of Science Core Collection (Clarivate Analytics, United States) was chosen as the data source. In March 2023, we conducted a search of the topic (phrases appearing in titles, abstracts and keywords) using the following search terms, “Sceletium” OR “mesembrine” NOT “Gastropoda”. A bibliometric data analysis, for the period 1961-2023, was used to determine trends within previous investigations and how *Sceletium* research has evolved, through tracking patterns, trends, relationships and the development of a discipline over time. Titles and abstracts were screened to exclude false-positives (papers that were not exclusively on *Sceletium* or mesembrine-type compounds found within the *Sceletium* genus). No supplementary restrictions had been placed on document type (review, editorial, letter, etc.) and assay model (*in vivo*, *in silico*, *in vitro*, etc.). The average citation amongst the most popular thematic areas within the body of knowledge associated with *Sceletium* is represented as a bar graph generated in Excel.

The data from our WOS searches were read from a bibliographic database file (i.e. the .txt file). Different types of analyses were performed based on our research questions. We were interested in determining the following: 1) the number of contributions in the field and how this changed with time; 2) authorship patterns; 3) geographical location of the producers of the articles, and finally; 4) identification of trends and gaps in the field.

#### Term maps

Term maps were generated using words in the titles and abstracts whilst authorship and country maps were generated from information associated with the authors and affiliations. Within the bibliographic analysis, 296 articles were analysed and visualized by VOSviewer (Van Eck and Waltman, 2010). VOSviewer is a software that visualizes patterns between authors, countries and terms found in a body of literature. The software is able to create networks between data and illustrate them as bubbles connected by lines, indicating association. The larger the bubble the greater its frequency of occurrence. The thicker the lines the greater number of links an item has with others in the network. Irrelevant phrases or repetitions of phrases were excluded.

## Discussion

### Past and current trends in literature

From the 296 articles that were published on *Sceletium* and Mesembrine-type alkaloids from *Sceletium*, the document types were predominantly articles (n=243) and reviews (n=46). The remaining major document types were scientific conference and meeting abstracts (n=4) and proceedings papers (n=3). The citations received by the 296 articles in this domain ranged from 0 to 208 (mean ± SD=26.02 ± 28.18). The most cited paper was between two papers, the first an ethnobotanical review by Stafford et al. (2008) investigating traditional South African plants with CNS activity (8.86 citations per year). This was followed by the Gu and You, (2011) paper on the organic synthesis of mesembrine isomers (11.27 citations per year). The hundred most cited papers within the field had an average citation of 55, with an average yearly citation of 4.

The thematic areas where the majority of the research is focussed are: Chemistry; Molecular Biology; and, Pharmacology. The average citation amongst the most popular thematic areas associated with *Sceletium* research is represented by Fig 4. For this reason, this review has a stronger emphasis on the work conducted in these fields. A particular focus has been placed on one species, *S. tortuosum* (119 links to other topics), and the membrane-rich extracts (64 links to other topics) of this plant. This has been the trend since the initial scientific interest in the plant in the 1960s. It is also interesting to note a lack of publications between 1980 and 2000. Dominant investigation areas were identified as ‘chemistry’ and ‘pharmacology’ especially, those focusing on *Sceletium* alkaloids to further understand the medicinal application of this plant (Fig. 5).

**Fig. 4.**
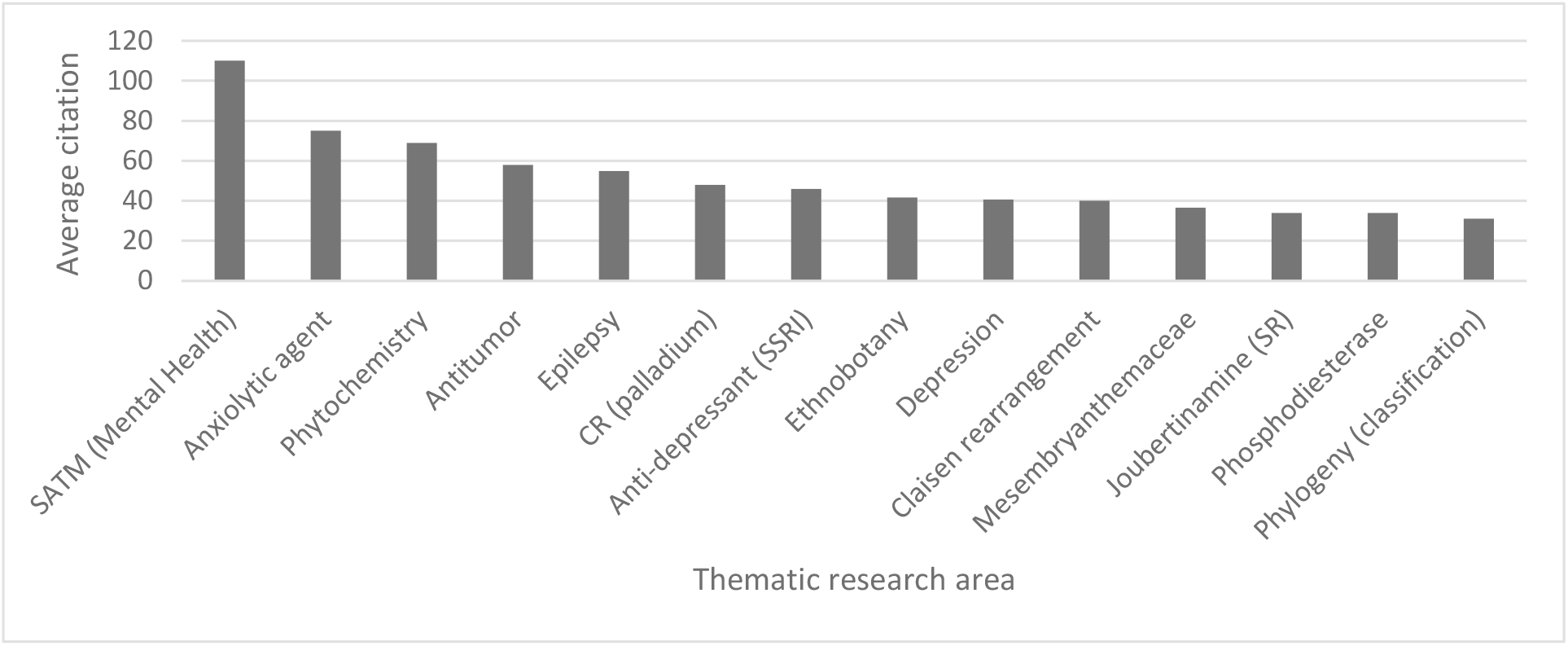
Average citation of thematic research areas within *Sceletium* research from Web of Science (n=296, citation values>30.00). SATM (Mental health) = South African Traditional Medicine (Mental Health); CR (palladium) = Coupling reactions (palladium); Joubertiamine (SR) = Joubertinamine (sigmatropic rearrangements)

**Fig. 5.**
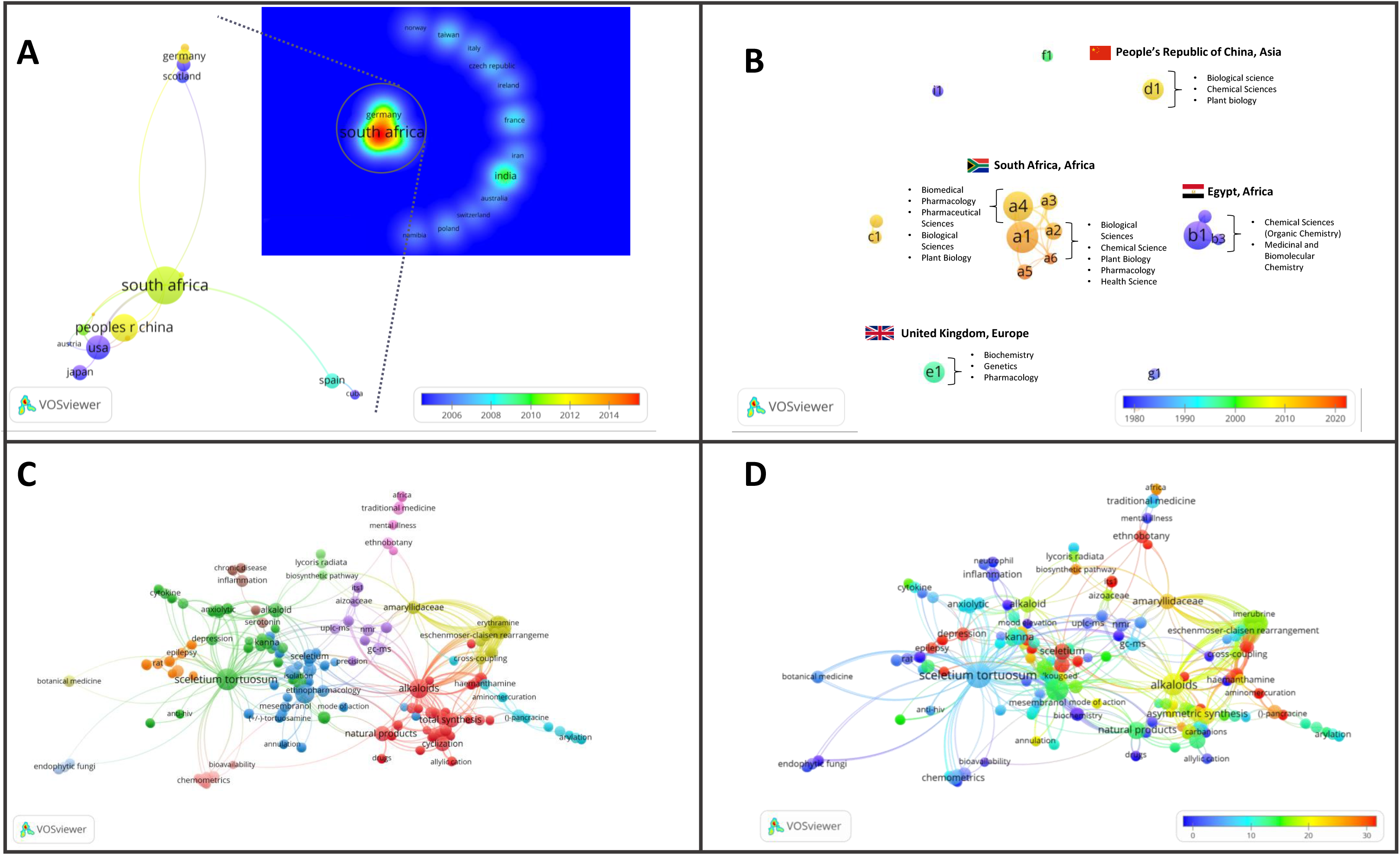
Major bibliographic summaries of literature in *Sceletium* research. a: Country network map of the most prolific research network in *Sceletium* research, based on country affiliation. Image additionally illustrating Average publication year; b: Author network map showing linkages and collaboration between various researchers (and institutions) with an overlay of Average publication years weighted by citations. (Map illustrating authors with at least 4 publications in the area of research).With overlay of associated countries and thematic areas.; c: Term map based on co-occurrence of text in both the title and abstract fields using 212 publications based on *Sceletium* research. d: Term map of research related to Sceletium with an overlay of the trend in citations over time. Data extracted from Web of Science (n=212)

### Key research themes

Several different research themes appear to be of superior relevance (as indicated by citation trends) in *Sceletium* literature. There were 349 terms that occurred three or more times in the 296 articles (Fig. 5 block D), these were separated into 13 thematic clusters identified through the VOSviewer (Fig. 5 block C). An analysis of the citations from 1956 to 2020 suggests that research associated with neurological disorders (ageing, depression, and anxiety) received significantly more citations per article (110, 55 and 41 average citations respectively). This can be seen by the red-colored bubbles (Fig. 5 block D). The neurological topics of ageing, anxiety and depression had an average of 110, 75 and 40.7 citations each, respectively. Other topics that were relatively highly cited were terms associated with the chemical synthesis of mesembrine alkaloids. These terms, C-H-amination, Claisen rearrangement, cobalt catalysis and enantiospecific synthesis had average citation values of 45, 40, 31 and 58 respectively.

### Drivers of research

The drivers of research in terms of authors came from 17 authors who had contributed findings associated with *Sceletium* and mesembrine (Fig 5B). These authors were selected on the basis of contributing 4 or more publications from 1956 to 2023. Among the authors, 3 networks can be observed. The network from South Africa is the greatest contributor in terms of publications, with the leading author contributing 15 papers on the topic. This may be due to their location which allows ease-of-access to wild-growing plant materials and established working laboratory methods, where plant material is sourced through permits for collection that is not destructive. The network from Egypt contributed 13 documents on the topic. Presently, within South Africa, the Tshwane University of Technology (averaging 11 citations per year) and Stellenbosch University (averaging on 10 citations per year) are the major contributors to research and have contributed the most publications with 18 and 10 publications respectively.

In terms of highly cited papers, with regard to chemistry from 1967 to 2000 the focus was largely on the isolation and characterisation of alkaloids from *Sceletium* species. Post-2000, the focus shifted to more chemical assays in an effort to develop quality control tools for the medicinally important plant, *S. tortuosum*, that was gaining pharmacological traction in literature as a phytomedicine used for anxiety, depression and as a mental stimulant. VOSviewer maps and an analysis of the literature indicates that the field may be shifting towards a greater focus on the toxicological and pharmacodynamics aspects of these plants (Fig. 5C). Gaps in the field of pharmacology in the field were identified as clinical trials and bioavailability studies.

We observed 29 countries/territories with the highest contributing countries being South Africa (50 documents), China (29 documents) and the USA (23 documents) where scientific investigations in *Sceletium* have been conducted. All three are associated in a network, and as such, have exchanged techniques and gained access to analytical tools for more advanced chemical and pharmacological analysis. South Africa is suspected to be the greatest contributor to current research efforts and this may be due to South African researchers having easier access to plant materials that grow in remote locations in the country and their compliance with Biodiversity laws that govern the issue of collection permits and bioprospecting activities linked to indigenous and endemic plant species in the country. Outside of this collaboration network, India has contributed 14 documents without collaboration with South Africa and these documents mainly cover topics related to the synthesis of mesembrine and joubertiamine alkaloids from *Sceletium* and one paper on the quality control of medicinal plants (Kumar and Sharma, 2018).

### Analytical Chemistry

The bibliometric analysis, performed in VOSviewer, identified the frequency and associations of keywords (Fig. 5c) and thus illustrates that alkaloid chemistry and analytical techniques have been a major area of interest for investigations linked to *Sceletium*. This is seen as the most dominant cluster in terms of publications, indicated by a red cluster. The most commonly used mesembrine alkaloid biomarkers in quality control and analysis are mesembrine (Fig. 2a), mesembrenone (Fig 2b), mesembranol (Fig 6, compound 2) and mesembrenol (Fig 6, compound 3). Some alkaloid classes which have been under-represented in the literature that may hold medicinal activity but have not been tested yet are joubertiamine (Fig 6, compound 5), sceletium alkaloid A_4_ (Fig 6, compound 6) and the tortuosamine alkaloid classes of compounds (Fig 6, compound 8), that are also found in *Sceletium* species. The purple cluster represents experimentation related to the isolation and identification of compounds by various analytical means (Fig. 5c). The body of work is quite substantial and has had a wide range of analytical techniques applied to the phytochemical characterisation of *Sceletium* species and related commercial products (Table 1). The main analytical techniques that seem to be predominantly applied include Ultra Performance Liquid Chromatography - Mass Spectrometer (UPLC-MS), High Performance Liquid Chromatography - Mass Spectrometer (HPLC-MS), Nuclear Magnetic Resonance (NMR) and Gas Chromatography–Mass Spectrometry (GC-MS). The majority of analytical methods used on *S. tortuosum* have targeted the detection of two alkaloids, mesembrine and mesembrenone (Table 1).

**Fig. 6.**
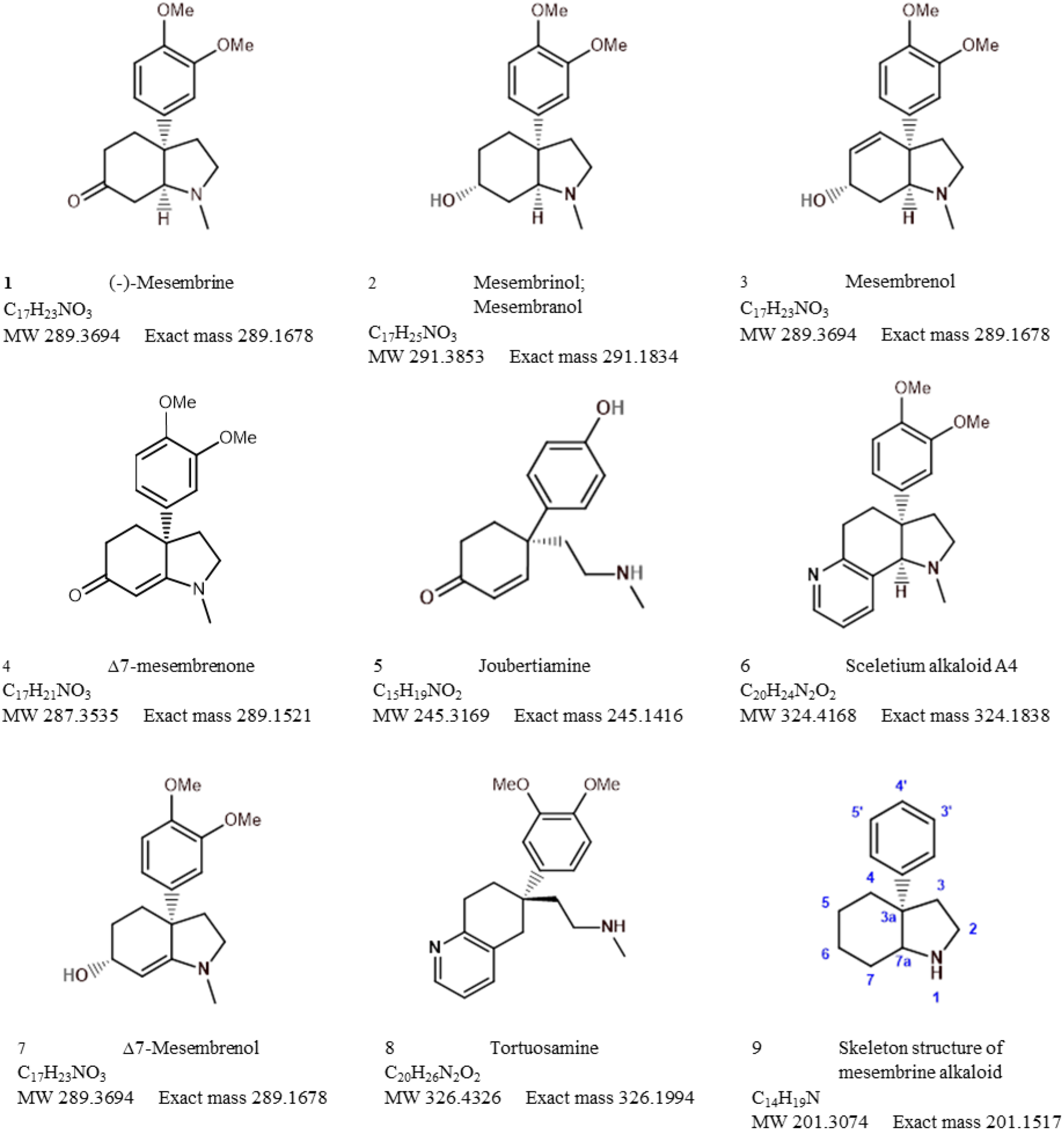
Mesembrine alkaloids from Sceletium of notable medicinal activity

**Table 1:**
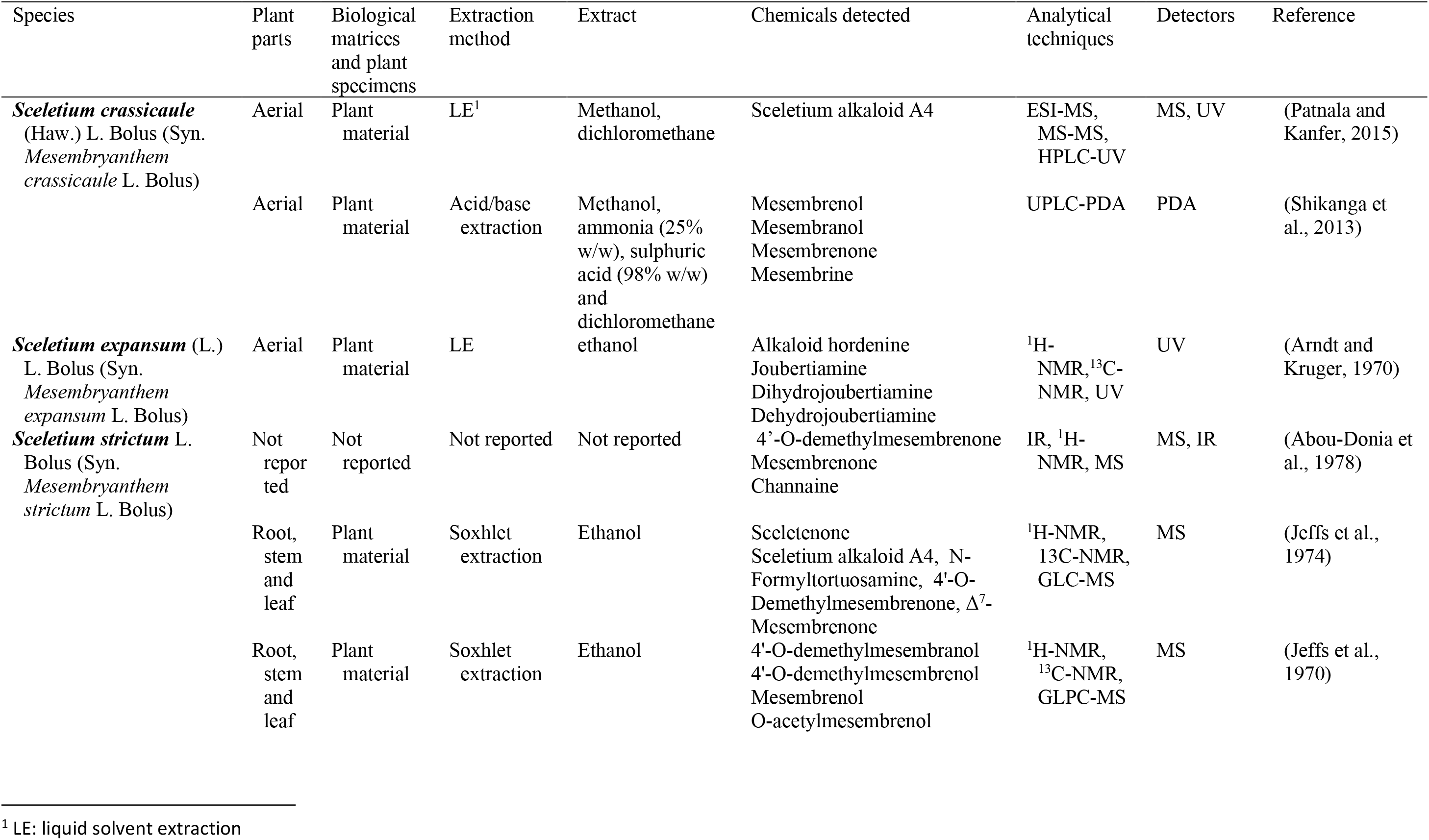
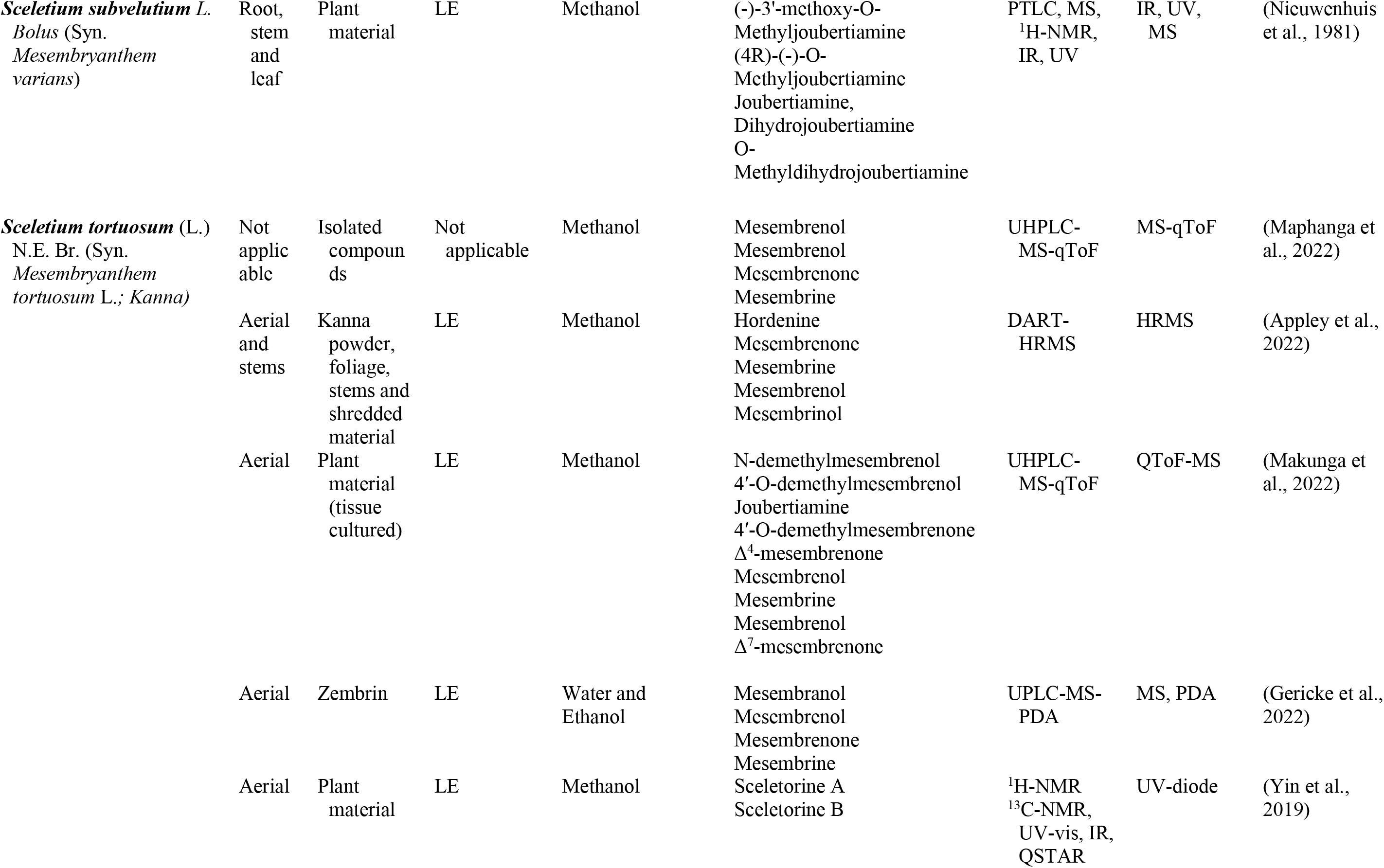
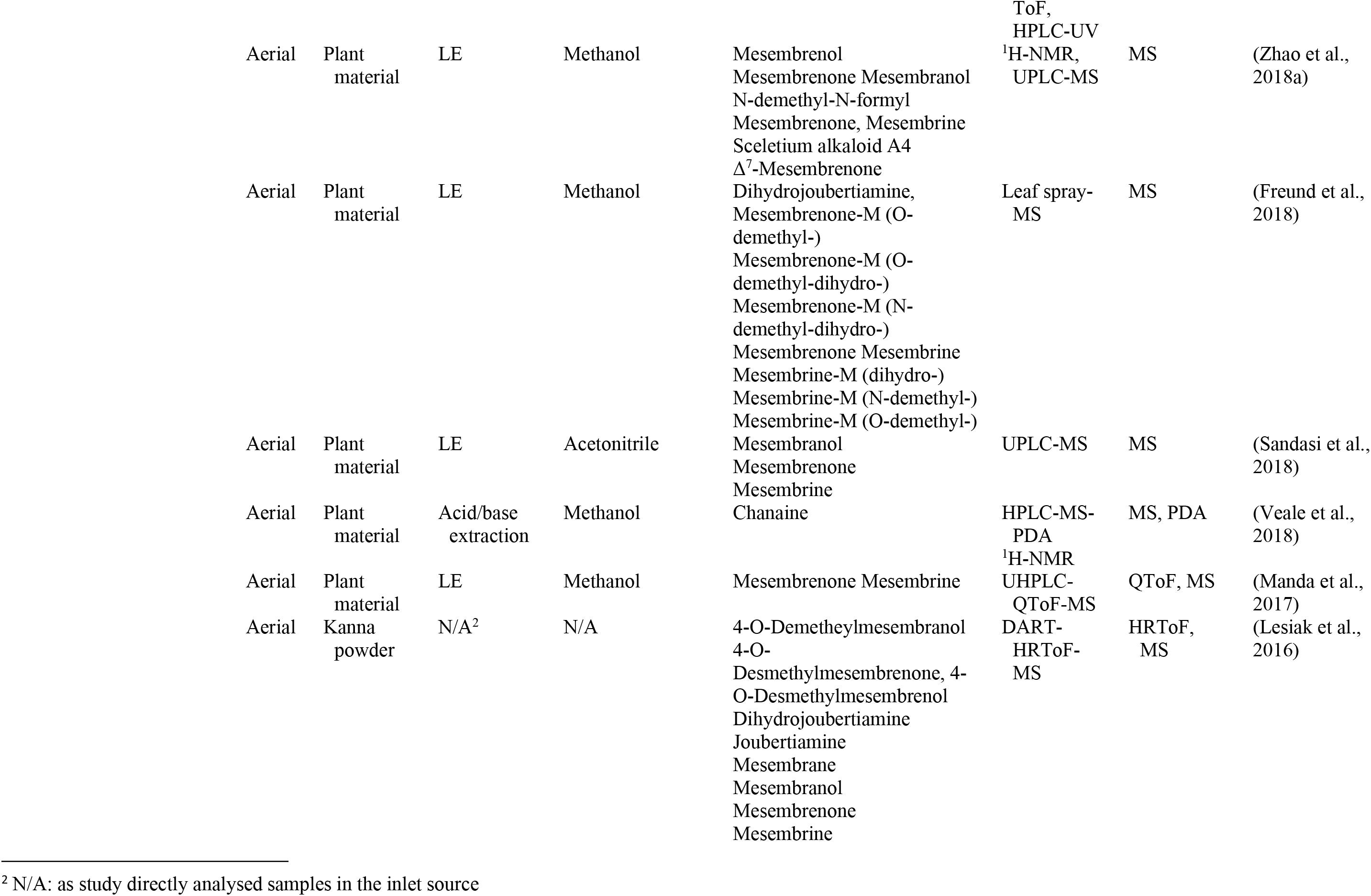
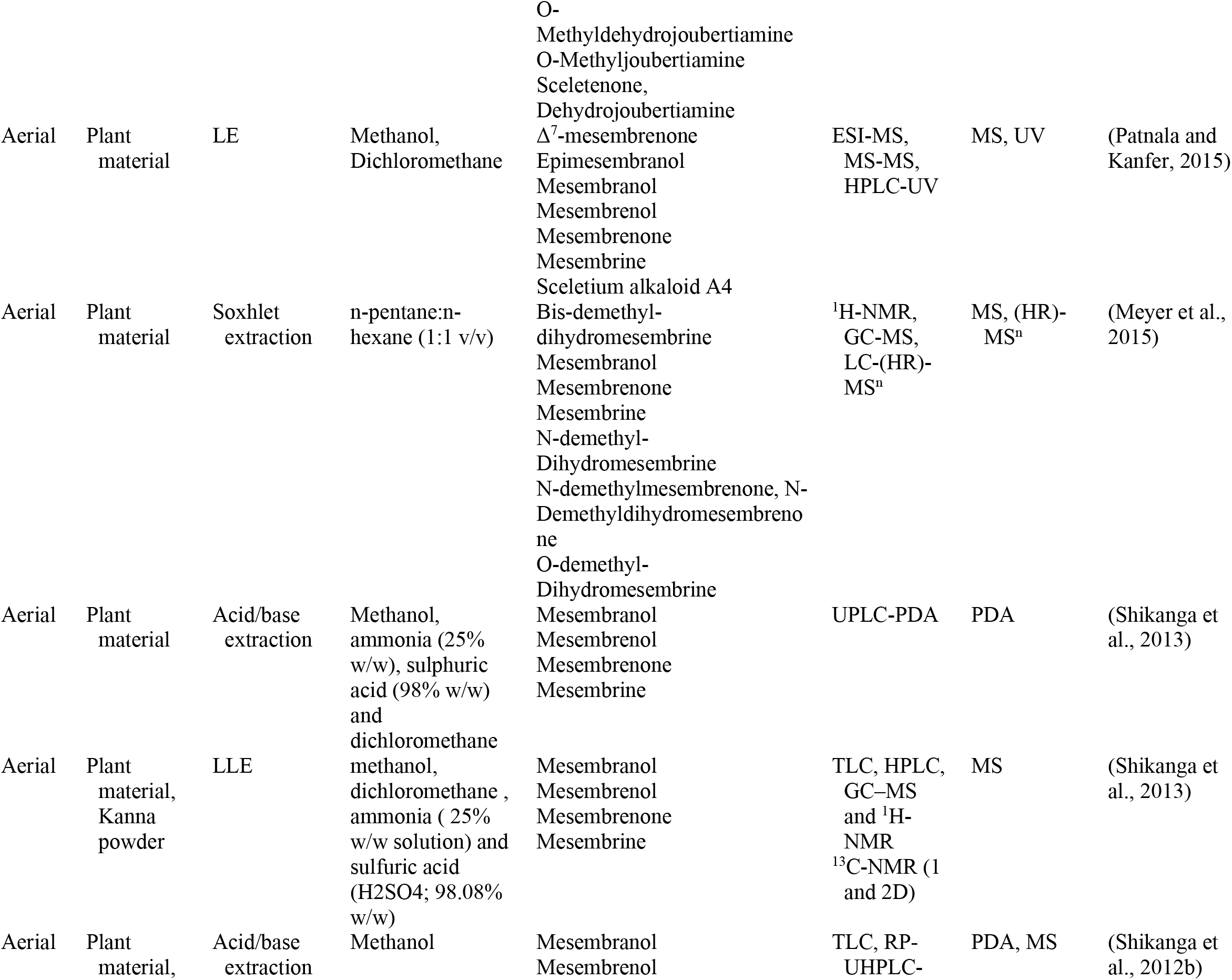
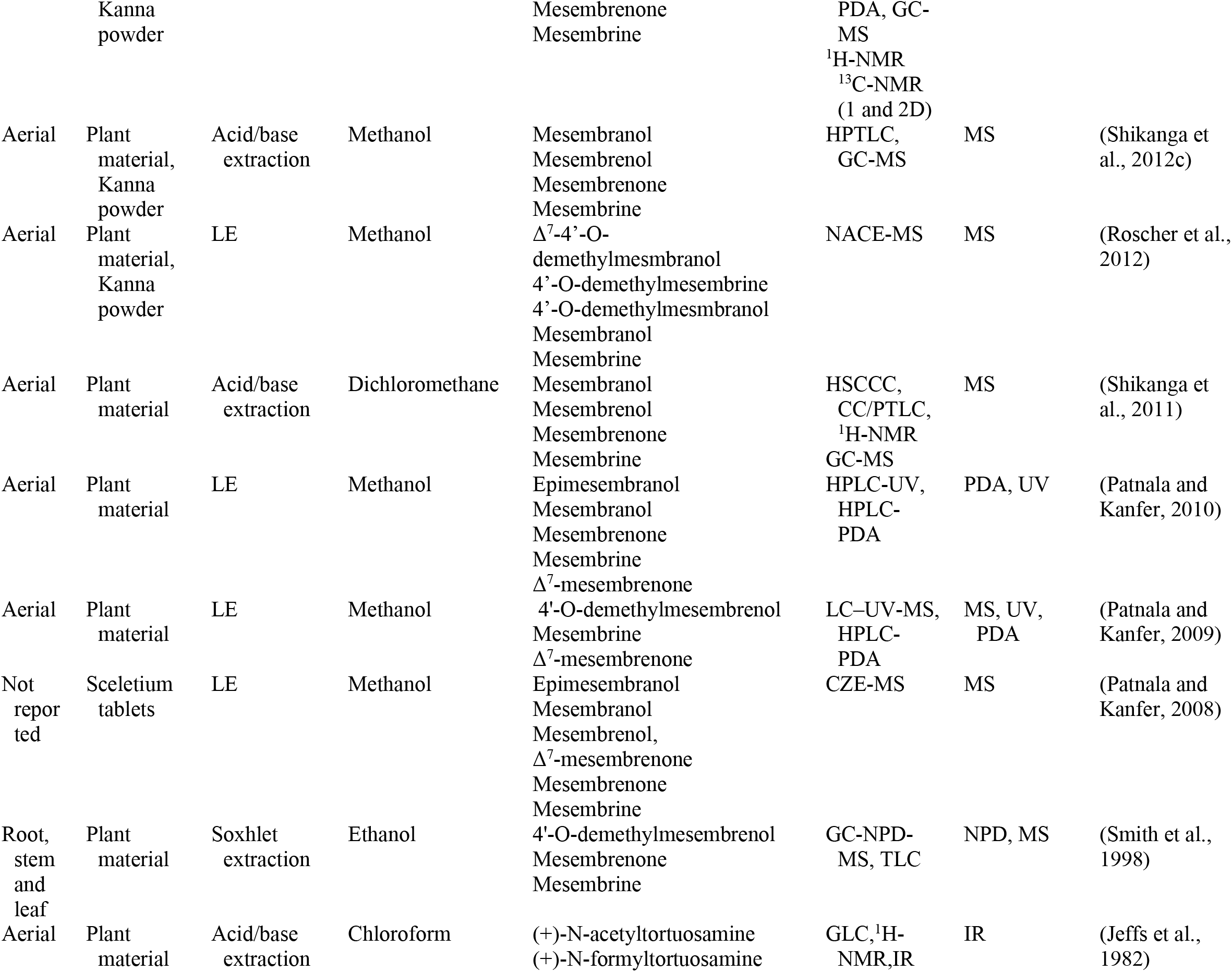
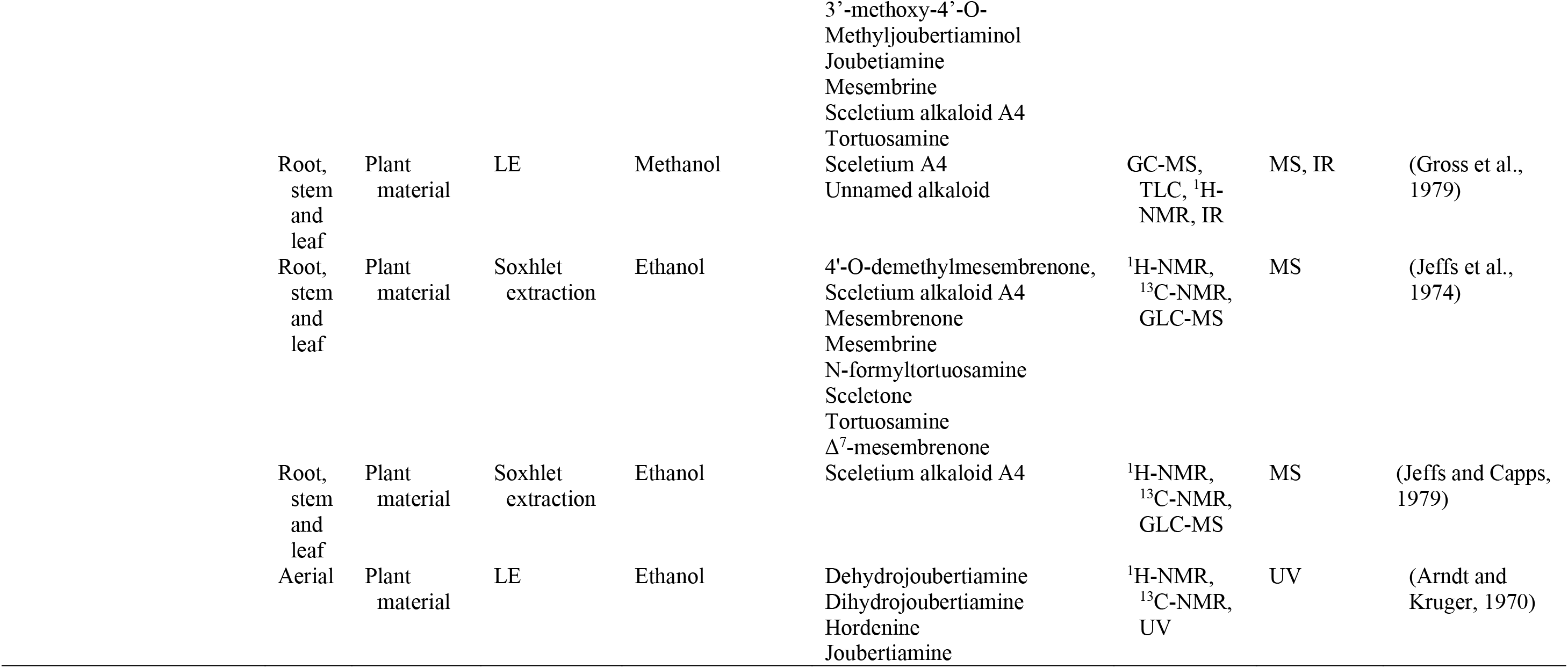
The analytical techniques used on Sceletium species of medicinal importance, with their extraction method, sample preparation and detectors (LE=Liquid Extraction)

Analytical chemistry in *Sceletium* has been of scientific interest since the 1970s. Many of the studies in the 1970s were mainly focused on compound isolation and structural elucidation (Jeffs et al., 1970, 1974b, 1974a; Arndt and Kruger, 1971; Abou-Donia et al., 1978; Nieuwenhuis et al., 1981), as examples). For these purposes, large amounts of material are needed to obtain metabolites in plants that generally occur in small quantities. This may largely have been driven by the search for novel phytochemicals that would be of interest for pharmaceutical companies, at that particular time. In terms of *Sceletium*, however, the isolated structures did not necessarily enter into a drug discovery pipeline during the period of 1970 to 1998. The analysis of crude extracts using a variety of techniques, from thin layer chromatography, gas chromatography and liquid chromatography is more evident in the literature and various laboratories have published several papers that focus on analyzing mesembrine alkaloids. With changes in research foci in the natural products industry, where the study of complex plant mixtures using metabolomics in the 2000s till present has become an established field, smaller quantities of plant materials are being utilised. Secondly, the focus has shifted to using such strategies for quality assessment of wild-harvested *Sceletium* species, as a means to compare wild populations to define chemotypes that occur naturally. Also, such application of metabolomics is explored for its potential contribution to the development of quality assurance protocols to ensure that *Sceletium*-based products are scientifically verified to contain the biomarker mesembrine alkaloids, that define their biochemical makeup (Masondo and Makunga, 2019). In a chronological format, the different analytical methods that have been used in *Sceletium* phytochemical studies are discussed below.

Jeffs et al. (1970), performed an ethanol extraction on *S. strictum* plant material. Plant material was ground, soaked overnight in 95% ethanol and then filtered *in vacuo*. The extraction of alkaloids was achieved by acid-base extraction, yielding 4.02 g of crude alkaloids (2.6% of dried weight). Gas-liquid partition chromatography (GLPC) was used for the determination of mesembrine type alkaloids in the crude extract. Two columns were used in this analysis, namely, SE-30 and Carbowax 20M. Both were glass columns containing 3% SE 30 on Aeropak 30 (100-120 mesh) at column temperature 220°C and 4% Carbowax 20M on Aeropak 30 (100-120 mesh) at column temperature 250°C, respectively. The crude alkaloid fraction was then analysed by chromatography over neutral alumina (200g) with a linear phase of benzene (2L) against ethyl acetate (2L), 200 mL ethylacetate-C_2_H_6_OH (4:1) and finally 500 mL of CH_3_OH. The GLPC allowed for the collection of 7 mg of an unidentified compound, 32 mg mesembrenone, 13 mg mixture of mesembrine-mesembrenone, 285 mg mesembrenol, 871 mg mesembrenol-mesembranol (at a ratio of 90:10 (w/w) and 101mg mesembrine. All of the structural identifications were possible and were achieved after nuclear magnetic resonance (NMR).

Another study by Arndt and Kruger, (1970), reported on an extraction of *S. joubertii* (syn. *S. tortuosum*). Contrary to the work of Jeffs et al. (1970), they reported that it was the first *Sceletium* species that did not contain mesembrine. Extraction of aerial parts (3.30 kg) of the plant was performed using 2% ethanolic tartaric acid, which yielded a crude alkaloid mixture of 1.01 g. Separation of the alkaloids was achieved by chromatography on an alumina column with Na_2_CO_3_. Identification of the compounds was achieved by NMR. Alkaloids identified in the material were mesembrane, joubertiamine, dihydrojoubertiamine, dehydrojoubertiamine and hordenine. The quantities were not determined for these compounds. Disputes in terms of the presence or absence of mesembrine in *Sceletium* species continue to date and this is largely due to differential accumulation of mesembrine alkaloids in plant material that may be collected at different times and/or localities. Misidentification of plant material can also occur and not all the species have the same profile of mesembrine alkaloids. It is thus important to report on the provenance of plant material and some of the early works with this particular plant did not necessarily take this into great consideration.

Sceletium A_4_ is one of the compounds that is now regarded as one of the biomarker compounds of *S. tortuosum* and together with mesembrine, mesembrenone mesembranol and mesembrenol (Figure 2), it can be used to differentiate wild-harvested chemotypes (Masondo and Makunga, 2019). The paper of Jeffs et al. (1971b), identified this particular compound, Sceletium A_4_ from *S. namaquense* (syn. *S. tortuosum*), for the first time. However, they did not report on any chromatographic techniques or quantitative data and information on structural characteristics from NMR data was solely presented.

N-demethylmesembrenol and N-demethylmesembranol were first isolated using 2% ethanolic tartaric acid and column chromatography (Arndt and Kruger, 1971). A crude alkaloid mixture (6.0g) was separated on a silica gel column with Na_2_CO_3_ using methylene chloride-methanol as a solvent system. Although quantification of this extract was not performed, NMR enabled the structural characterisation of the chemical isolates.

Jeffs et al. (1974) identified the alkaloids Sceletium alkaloid A_4_, tortuosamine, N-formyltortuosamine and sceletenone isolated from *S. namaquense* (syn. *S. tortuosum*). The isolation of alkaloids from *S. namaquense* was conducted using a Soxhlet extraction where 3.5 kg of plant material was extracted with 15 L of 95% ethanol for 17 hr. The material was then macerated with 10 L of 95% ethanol and an acid-base extraction was performed. Further purification of sceletenone alkaloid fraction was subjected to high-pressure chromatography on phenylcorasil in a water:acetonitrile solution (9:1; v/v). The authors report on the generation of several different fractions and the quantification of fractions from *S. namaquense* gave 50 mg tortuosamine, 94 mg sceletium alkaloid A_4_, 40 mg N-formyltortuosamine, 94 mg sceletenone, 95 mg mesembrenone and 40 mg Δ^7^mesembrenone. For this particular study, two different techniques, ^1^H-NMR spectroscopy and IR spectra, were used to confirm structural identities. The structurally similar mesembrine alkaloids (mesembranol, mesembrenol, mesembrenone and mesembrine) all contain the cis-3a-aryloctahydroindole nucleus (Krstenansky, 2017). This is advantageous as the utilisation of different techniques to confirm the structural composition of phytochemicals provides better resolution. Using X-ray crystallography, Abou-Donia et al. (1978) identified the structure channaine from *S. strictum*. However, the authors cautioned and suspected that channaine may have been an artefact from the condensation of two *N*-demethylmesembrenone molecules during the isolation process. With the exception of the recent study by Veale et al. (2018), discussed below in detail, no other reports have shown this unusual alkaloid to occur in *Sceletium* plants since then.

Nieuwenhuis et al. (1981), isolated O-methyljoubertiamine (16 mg) and O-methyldihydrojoubertiamine (10 mg) from *S. subvelutium* (syn. *S. varians*). Plant material (5.7 kg, whole wet plant) was extracted by homogenizing with 2% methanolic tartaric acid solution (5 L). The solution was left to stand for 12 days and filtered through celite. After which it was re-extracted with boiling methanol, left overnight and then filtered. An acid-base extraction was performed, followed by liquid-liquid partitioning. The extract was separated in a chromatography column on basic alumina (25 g, packed with anhydrous benzene). The column was eluted with benzene (75 ml), benzene-chloroform (4:1, v/v, 50 ml), chloroform-benzene (4: 1, v/v, 50 ml), chloroform (20 ml), chloroform-methanol (1:1, v/v, 50 ml) and methanol (75 ml). Dragendorff’s solution was sprayed at the end for the identification of alkaloids and finally, compounds were confirmed by NMR. Another report by Jeffs, Capps and Redfearn (Jeffs et al., 1982), performed an extraction on *S. namaquense* (syn. *S. tortuosum*). In this particular study, 3.5 kg was used as the starter material for the extraction with 15 L of 95% ethanol being used as the extractant for 17 h. The material was then macerated with 10 L of 95% ethanol and an acid-base extraction was performed before the mixture was filtered. The mixture was fractionated in a chromatographic column with 1200 g neutral alumina before the column was eluted with 3 L of solvent in a CHCl_3_-CHCl_3_/MeOH (4:1) gradient combination. The isolated compounds obtained were: 1) non-alkaloidal material that weighed 3.70 g; 2) mesembrine recorded at 1.63 g; 3) 1.5 g mesembrenone, 4) Sceletium A_4_, (-) -3’-methoxy-4’-0-methyljoubertiamine, and, other alkaloids all being recorded at a mass of 2.5 g; 5) 4-(3,4-dimethoxyphenyl)-4-[2-(acetylmethylamino)ethyl]cyclohexanone and unidentified alkaloids are reported as weighing 1.45 g. The other phytochemicals that were isolated were N-formyltortuosamine and unidentified alkaloids (3.11 g) together with Δ^7^mesembrenone and tortuosamine being recorded at 1.11 g. The authors again utilised an NMR-based method for the structural identification of the above-mentioned chemical isolates.

There is a clear lack of analytical isolation methods being used in studying the chemical constituents of *S. tortuosum* and its relatives from the 1980s till 1998. Renewed interest in these species is evident thereafter, with the work of Smith et al. (1998) that tested 21 species from 9 genera of the Mesembryanthemaceae, for the distribution of mesembrine alkaloids. As compared with previous studies, significantly less plant material was used per extraction with methods staying relatively the same. The analytical techniques had advanced quite notably with the last research on *Sceletium* and its alkaloids that had been performed 16 years prior. Many investigations after 1998, use smaller amounts of the plant sample as techniques that are in routine use for metabolite profiling are much more robust. Alkaloids were extracted from root, stem and leaf tissues using 25 g each of each plant part and a total of 75 g of being material thus being used. A soxhlet extraction with 95% ethanol for 12 h was performed. This extraction was followed by separation in a column packed with 60 ml Extrelut (a specially processed, inert, wide pore Kieselguhr with a high pore volume). The solvent system used in the column was 40 ml dichloromethane: isopropanol (85:15; v/v). The solvent was removed and samples were run using column chromatography again with the following solvents: dichloromethane, ethyl acetate, acetone, acetonitrile, methanol and acetic acid. These fractions were run on TLC plates before Dragendorff’s reagent was sprayed on the plates to confirm the presence of alkaloids. The studied fractions were concentrated and analysed using gas chromatography (GC) fitted with a nitrogen–phosphorus detector (NPD). The column heating program was run from 230°C to 260°C at a rate of 1°C per min. The detector was set at 350°C. The GC-MS unit was fitted with a 30 m fused silica DB-5 capillary column of 0.25 mm internal diameter before 1 µl samples were injected into the GC/MS using the same column temperature program as for the GC/NPD work. For reproducibility and to correct for day-to-day variation in retention times on the NPD detector, a *Sceletium* extract sample was run between and after samples. Only semi-quantitative data could be generated for this study due to no standards being readily available at that particular time. A mass spectrometry (MS) method was employed in order to determine several different alkaloids with retention times and analysed chemical isolates were as follows: 4’-O-demethylmesembrenol (11.5 min; [M]^+^ 1 275 (55%), [M-1]^+^ 1 274 (33%), [M-Me]^+^ 1 260 (6%), 232 (15%), 219 (28%), 218 (35%), 204 (100%), 137 (9%), 110 (8%), 96 (92%), 70 (98%)), mesembrine (12 min; [M] ^+^ 1 289 (66%), [M-1]^+^ 1 288 (36%), [M-Me]^+^ 1 274 (9%), 246 (8%), 232 (19%), 219 (48%) 218 (92%), 204 (48%), 151 (11%), 110 (4%), 96 (100%), 70 (87%)) and mesembrenone (12.5 min; [M]^+^ 1 287 (21%), [M-1] ^+^ 1 286 (4%), [M-Me]^+^ 1 272 (2%), 259 (4%), 258 (5%), 244 (2%), 230 (3%), 219 (7%), 218 (2%), 204 (2%), 149 (2%), 110 (, 1%), 96 (2%), 70 (100%). Out of the species tested, the only species with comparable mesembrine alkaloid levels to that of the *S. tortuosum* was *Aptenia cordifolia*. The relative levels of alkaloids as a percentage of the mesembrine pool in *Sceletium* were 0.3 (7.2 min), 0.2 (8.7 min), 4.6 (9.2 min), 14.4 (11.5 min) and 9.7 (12 min), respectively. The relative levels of mesembrine were not reported on nor were *m/z* data for the other species presented.

Methods that allow for high-throughput detection of mesembrine alkaloids are thus sought after for industrial applications. Such methods also need to be less labour intensive and not necessarily require a high level of technical know-how for them to be placed in routine use, more especially to use them as a quality assurance measure and for the standardisation of manufactured products derived from *Sceletium*. As an example; Patnala and Kanfer, (2008) investigated the analytical technique of capillary electrophoresis (CE) as a possible method to study isolated phytochemicals from *Sceletium* as well as commercially available *Sceletium* supplements. This is a technique where electrophoretic mobilities under the influence of an applied electric field enable the separation of charged components. This analytical technique allows for the rapid and efficient separation of compounds leading to rapid analysis (Li, 1992). The technique is favourable due to its high efficacy and efficiency, wide application for both scientific laboratories and industrial manufacturers plus it requires low running costs during experimentation. Major compounds and biomarkers were extracted from *Sceletium*, namely, mesembrine, mesembrenone and Δ^7^mesembrenone and two other compounds, mesembranol and epimesembranol, were produced by performing a hydrogenation on a pure chemical isolate of mesembrine. The CE analysis was paired with a diode array detector (DAD) and a UV detector before NMR analysis was also performed for the characterisation of phytochemicals. Separation of individual alkaloids was carried out by 50 cm effective length, fused silica capillary tubing (50m i.d. × 360m o.d.). Capillaries were conditioned with 1 M NaOH solution for 30 min, 0.1 M NaOH solution for the next 30 min followed by water for 40 min, with pressure of > 2500 mbar. Washing of the capillaries between consecutive injections to ensure optimal charge density on the capillary wall was performed with water (4 min), 1 M NaOH (2 min), 0.1 M NaOH (2 min) and a final washing step with water for 5 min was included in the method. Conditions during analysis were: ambient temperature of 22 ± 2 °C, applied voltage of +16 kV with a voltage ramp of +6 kV/s and background electrolyte (BGE) NaH_2_PO_4_·2H_2_O (50 mM). For this particular study, the pH was adjusted to 1.5 with H_3_PO_4_. Before this study, there was a paucity of reports on commercialised products of *S. tortuosum* despite an industry that had become established in South Africa. The study found the average content of mesembrine per tablet to be 164.30 μg per 12 mg dose of a tablet. Sensitivity and reproducibility were also an important consideration and the authors confirmed that their protocol was both sensitive and reproducible. They did this through an analysis of inter-day variation over 3 days and the precision recorded for mesembrine, as a quality control standard, was less than 4%. Importantly, the limits of detection (1.5 g/ml) and quantitation (2.5 g/ml) of the method were also reported by the authors. However, the exact species of *Sceletium* is not reported on in this study which hinders the reproducibility of this work as one can merely assume that the focus was on the commercialised plant, *S. tortuosum*. It should be noted that there were some encountered difficulties during experimentation as the method could not conclusively distinguish between compounds with similar *m/z* values (diastereomers at *m/z* 292). Correct taxonomic identities for *Sceletium* species need to be accurate as these plants are difficult to distinguish from their anatomical structures and chemotaxonomic markers have not always played a significant role in delineating sister species from each other (Patnala and Kanfer, 2013).

Traditionally, *S. tortuosum* is claimed to have been fermented before it was used as a euphoric agent in the spiritual rituals of the Khoe-Sān. Such a practice points to the importance of fermentation. In some instances, those that farm this product prefer to ferment the material as they believe that it may increase the biomarker compounds that are assessed in quality assurance protocols associated with the *S. tortuosum* herbal industry (Olivier L, personal communication). Several analytical studies have thus aimed at determining the phytochemical content of *S. tortuosum* and the effect of fermentation on its profile. The techniques to study fermented plant materials of *Sceletium* are varied in their approach.

The analytical tools chosen by Patnala and Kanfer, (2009) were HPLC-PDA, LC-MS with a UV detector for qualitative and quantitative analysis of 75g of aerial material which was subjected to bruising, fermentation and extraction with methanol. Two methods of fermentation were used in their study. The first involved crushing 75 g of the aerial parts in a plastic bag before the material was left to ferment in the sun. Samples were then removed and analysed from the bag daily over the course of 10 days. The second analysis of the fermented materials was conducted a year later using 130 g of plant material. Fermentation resulted in the transformation of mesembrine. This subsequently led to lower levels being detected which confounded evidence presented by Smith et al. (1998). Although precision and sensitivity were reported on, the authors did not provide detailed information relating to instrument conditions, making this study rather difficult to reproduce in other laboratory environments.

Apart from their interest in the fermented profiles of *S. tortuosum*, Patnala and Kanfer, (2010) developed and validated an HPLC method for the analysis of *Sceletium* plant material but the exact species used for this investigation was indicated and one assumes that *S. tortuosum* was the target species. The method showed repeatable, precise, and appropriate resolution of alkaloids for quality control of mesembrine-type alkaloids. Prior to this study, poor validation data had been presented on any analytical techniques described for *Sceletium.* The analytical tools used were an HPLC system connected to a UV and PDA detector. Further structural data was supported by NMR spectra. Two different columns were used in the HPLC system, namely, Luna® C_18_ (2), 5 μm, 150 mm x 4.6 mm i.d. and Hypersil® 150 x 4.6 mm i.d C_18_ column. The lack of standards resulted in several alkaloids assayed via chromatography whose identities were unknown to the researchers as no published data on these metabolites was available. The mobile phase in the column was water:acetonitrile:ammonium hydroxide solution (25%) mixed in a ratio of 70:30:0.01 (v:v:v) and was tested under isocratic conditions using the Hypersil® 150 x 4.6 mm i.d C_18_ column. This separated alkaloids but better separation (resolution) was achieved with a mobile phase of 0.1% ammonium solution in water combined with acetonitrile for 15 min on a. Luna® C18 (2) HPLC column. The alkaloids, mesembrine (RT 10.894 min), mesembrenone (RT 7.579 min) and Δ^7^mesembrenone (RT 5.242 min) were isolated and confirmed by NMR. The method was validated in terms of linearity, accuracy, recovery and limits of detection. Unfortunately, this study did not report on what species of *Sceletium* they had extracted the phytochemicals from, which is an issue in the reproducibility of this study. To iterate, this is highly problematic when some of the *Sceletium* species are difficult to distinguish from each other as they are similar in their appearance and their taxonomy is rather ambiguous (Patnala and Kanfer, 2013). The provenance of the plant samples can alter their phytochemical composition as many different chemotype configurations may exist in wild-collected populations, exhibiting both intra- and inter-specific variability (Shikanga et al., 2012d).

Due to the complex mixture of structurally similar alkaloids, the development of appropriate analytical techniques for chemotaxonomic assessment has proven to be quite a challenge. Further compounding this issue is that, in some species, for example, *S. emarcidum*, the distribution of alkaloids falls below the limit of quantification by the analytical tool (Patnala and Kanfer, 2013). To reduce these challenges, the introduction of reference compounds for all the alkaloids of interest may allow for better specificity during the fingerprinting process, assisting with the assay of plants with stronger precision and accuracy.

Shikanga et al., 2011) employed a high-speed countercurrent chromatography (HSCCC) method to rapidly isolate alkaloids from *S. tortuosum* in high yields. For the plant extraction 500 g of aerial parts of *S. tortuosum* were macerated in 0.25 M sulfuric acid (3 6 L), left to settle for 15 min and then filtered. The mixture was extracted again with dichloromethane (3 875 ml) and evaporated to dryness (9.68 g). The alkaloidal extract was then run through column chromatography (CC) set up, packed with 200 g Kieselgel 60. The solvent system used was CHCl_3_–MeOH gradient (100:0, 99.5:0.5, 99:1, 97:3, v/v). Fractions were analysed by TLC using the eluent system of CHCl_3_–MeOH–10% NH_3_ (90:10:0.1, v/v). Fractions were then analysed by HSCCC using a solvent combination of n-heptane–MeOH–EtOAc–1% NH3 (1:3:1:3, v/v). The machine was operated on normal mode at 30 °C, coils were rotated (1500 rpm) being pumped at 3.0 ml/ min (130 psi) until the mobile phase left the coils. For GC-MS analysis (determination of purity), 2 µl was injected at a temperature and pressure of 255 °C and 12.54 psi. The column used in the system was a HP-5MS 5% phenyl methyl siloxane column (30 m 250 mm i.d. 0.25 mm film thickness) with an oven temperature of 60 °C, rising to 255 °C at a rate of 20 °C/min and held for 15 min. The carrier gas used was Helium at a flow rate of 0.7 ml/min. compound purity was determined by integrated peak area. The alkaloid quantities (and purities) obtained by HSCCC were 482.4 mg (98.5%) mesembrine, 545.2 mg (98.4%) mesembrenone, 300.0 mg (98.2%) mesembrenol and 47.8mg (95.4%) mesembranol. The quantity and purity obtained by HSCCC was higher in all alkaloids as compared to CC/PTLC also performed in this study. The method was efficient and cost effective, requiring relatively smaller amounts of plant material in isolating mesembrine (482.4 mg), mesembrenone (545.2 mg), mesembrenol (300.0 mg) and mesembranol (47.8 mg).

Chemotypic variation observed in *Sceletium* (Roscher et al., 2012; Shikanga et al., 2012d; Zhao et al., 2018) may be due to the ability of plants to exhibit phenotypic plasticity to cope with their environments and climates (Nicotra et al., 2010). Phenotypic plasticity enables plants with a standard genome to adapt their phenotype in response to environmental pressures assisting with survival (Nicotra et al., 2010). This phenotypic plasticity is often correlated to metabolomic differences in the plants in response to their environments, an example of this was observed in *Hippophaë rhamnoides* (Kortesniemi et al., 2017). The study of chemical variation linked to plant-environment effects can thus easily be achieved using phytochemical analytical techniques.

The field of plant metabolomics is making major contributions to our understanding of plant biochemistry and metabolism as a metabolomics workflow can facilitate a comprehensive compilation of metabolites within a particular cell, tissue or organ but large-scale experiments are notoriously difficult to interpret. In such instances, the complexity of these data sets is enormous and they cannot easily be processed with classical statistics (Van der Kooy et al., 2008), consequently principal component analysis (PCA) and partial least squares (PLS) analysis have been employed. These types of multivariate statistical applications reduce the dimensionality of the data enabling better pattern recognition that can be correlated to the analysed samples. Shikanga et al. (2012d), performed a chemical analysis of different populations located at 12 different sites of *S. tortuosum* and analysed these samples by GC-MS to gain an understanding of chemotypic variation. The aerial parts of the plants were dried at 30 °C for two weeks after which material was pulverised, sonicated and extracted by an acid-base extraction method as reported by Alali et al. (2008) with slight modifications in the sulfuric acid used (0.5 M; 24.0 ml). Extracts were sonicated and vortexed, followed by transferring and neutralising the supernatant with 20% aqueous ammonia. The organic phase was separated by dichloromethane and evaporated to powder (the same was applied to commercial samples). For GC-MS analysis, 2 µl was injected into the system at a pressure setting of 24.0 psi and an inlet temperature of 250 °C. Initial oven temperature was kept constant at 259 °C for 2 min, then increased to 262 °C at a rate of 20 °C/min and held constant for 6 min. Separation was achieved using a HP-5MS 5% phenyl methyl siloxane column (30 m 250 mm i.d. 0.25 mm film thickness) and all runs used helium gas, maintained at a flow rate of 1.4 ml/min. Electron impact ionization was employed (70 eV; 35–550 m/z) and MS spectra were collected over a run time of 8 min. Total alkaloid yields across species were highly variable ranging from 0.11 to 1.99% of dry plant weight (dw). By using this approach, the authors could confirm the presence of different chemotypes that exist in nature and five chemotypes were identified in the different populations. These chemotypes were differentiated from each other based on four identified alkaloids which occurred at higher concentrations in the Calitzdorp populations with a total of 1.66% alkaloid as a yield. Purchased products were validated to be composed of several different mesembrine alkaloids with mesembrine occurring at a range of 42.26 to 82.94 %. This study showed the robustness of the GC-MS method in generating quality assurance standards for both wild-harvested plant materials and purchased products. In another study by Shikanga et al. (2012b), both GC-MS and ultra-performance liquid chromatography with photodiode array detector (RP-UHPLC PDA) were used as the metabolite profiling techniques but in this work, the acid-base extraction reported was compared to the use of methanol and water, that had previously tested in another study by Shikanga et al. (2011). The acid-based extraction was preferred as a method as it was reported to show better extraction of pyrrolizidine alkaloids that is known to have considerable toxic effects when consumed (Alali et al., 2008). However, no quantitative measurements were conducted for both water or methanol as extractants and so the authors make no mention of the quantities obtained when comparing these solvents. Details focus mainly on chromatographic aspects and the chromatography was achieved by a Waters Acquity reversed phase UHPLC BEH C_18_ (2.1× 150 mm, 1.7 μm particle size) column and a Van Guard pre-column (2.1× 5 mm, 1.7 μm). The analysis of samples in GC-MS was analysed using an Agilent 6890N gas chromatograph, coupled with a 5973 mass spectrometer. The column used in GC-MS was g a HP-5MS 5% phenyl methyl siloxane column (30 m × 250 μm i.d.× 0.25 μm film thickness). The method was validated in terms of linearity, limits of detection, repeatability and recovery. The method gave satisfactory separation of peaks (resolution) and quantification was possible in both analytical techniques. The analytes targeted for quantification were, mesembrenol (53.44 ± 1.15 μg.ml^−1^), mesembranol (25.23 ± 0.68 μg.ml^−1^), mesembrenone (35.42 ± 0.66 μg.ml^−1^), mesembrine (13.47 ± 0.30 μg.ml^−1^). Data obtained from both analytical methods were found to be similar. However, lower levels of alkaloids could be identified by RP-UHPLC PDA. The findings from this study indicate that the method is effective for the identification and quantification of four pharmacologically important alkaloids making these methods suitable for quality assurance procedures that may be employed during the production of phytopharmaceuticals based on *S. tortuosum*.

Shikanga et al. (2012c) further developed a method for the rapid and simple identification of alkaloids in *S. tortuosum* raw and wild-harvested materials. The intended purpose of this study was to develop an analytical technique for the routine analysis of psychoactive alkaloids in *S. tortuosum* these products. The analytical tool used was a high-performance thin-layer chromatography (HPTLC) densitometric method as this is a superior and more sophisticated form of TLC that is fast and robust for quality testing of botanical materials. One of the advantages of using this method is the automation of the different steps that would mainly be performed by hand with a normal TLC. This makes this method more powerful for metabolite fingerprinting, increasing its resolution and enabling quantitative measurement of phytochemicals. Because it is simple with many experiments being run at low costs, compared to GC-MS and/or LC-MS, it allows for rapid and parallel analysis of samples making it a high-throughput and reliable technology, that is now routinely used in many different settings for herbal drug authentication. As part of the protocol development steps, the HPTLC data of Shikanga et al. (2012c) were compared to a reference method that was previously developed in their laboratory based on GC-MS data of Shikanga et al. (2012b) using extracts generated with their acid-base method. Preparation and extraction of wild plant and commercial material (2 g per extraction) was performed as in Shikanga et al. (2012d). Sample application was performed using a 25 μL syringe and connected to nitrogen gas. Quantification was done by (UV)/vis densitometer, in absorbance mode at 280 nm. The method was validated in terms of linearity, accuracy, precision, and the upper and lower limits of detection. The HPTLC method was able to separate alkaloids, however, Rf values were quite close between mesembrenone (0.71) and mesembrine (0.60). Even so, peak purity was confirmed by UV-Vis. The repeatability of the method was confirmed by ANOVA but did not identify significant differences in analysis results between and within days (p < 0.05). Quantification of alkaloids in ng/band were, mesembrenol (275.09 ± 2.34), mesembranol 217.15 ± 3.40), mesembrenone (192.38 ± 2.27), mesembrine (160.09 ± 3.75). Data obtained from both analytical methods, HPTLC and GC-MS were found to be similar in terms of the power of detection. The HPTLC densitometric technique was found to effectively separate the biomarker mesembrine alkaloids from the other phytochemicals in the plant. This particular technique was thus recommended because it is simple and high-throughput in its application produces fast, reliable and reproducible results and can be routinely applied as an effective tool in the regular quality control of commercial and wild-harvested *S. tortuosum* samples.

The first study to use the analytical technique of non-aqueous capillary electrophoresis coupled to mass spectrometry (NACE-MS) for the separation of alkaloids in wild and commercial material was aimed at analysing wild and commercial plant materials was extracted using methanol as a solvent from *S. joubertii* (syn. *S. tortuosum*) (Roscher et al., 2012). The study aimed to analyse the alkaloid profile of *Sceletium* in an effort to provide a tool for the relative quantification of alkaloids in different *Sceletium* preparations. Another point of interest in this study was to investigate the influence fermentation would have on alkaloid profiles. Wild (calyx, stems and leaves) and commercial plant material was extracted using methanol as a solvent. Samples were also fermented and their alkaloid profiles were obtained. The fermentation was performed by crushing the whole wild harvested plant and this included the calyx, stems and leaves as plant parts. After homogenisation of the material, it was then stored in an airtight transparent plastic container, left in the sun for 8 days and vacuum dried later. All analyses were performed on capillary electrophoresis coupled to an Ion Trap 6330. The sheath liquid used in the capillaries (50 m id, length 70 cm) was 5% glacial acetic acid in isopropanol:water (66:34; v/v), at a flow rate of 4 μL/min. The method’s mean relative standard deviation for repeatability was 0.2% and 7.7%, respectively (n = 5) while the mean dynamic range was 0.1–500 μM. The CE separation was performed by applying 30 kV, resulting in a current of 16 μA. Identification of alkaloids was tentatively determined using mass fragmentation data and *m/z* values in comparison with previously published literature. The separation power of this technique is evident in its ability to separate fragments with similar *m/z* ratios, whereas the technique by Patnala and Kanfer, (2008) was unable to discriminate between two diastereomers of 4’-O-demethylmesembranol. The high selectivity of this method is evident by its ability to distinguish the diastereomers of 4’-O-demethylmesembranol (retention time of 12.3 min), since then no other analytical techniques have been able to identify it (Smith et al., 1998; Patnala and Kanfer, 2008, 2010). The NACE-MS method thus proved an effective method for separating isobaric structures in *Sceletium* samples with good resolving power. However, no quantification data were presented in this paper. Furthermore, no comparison is made with a reference technique with the same samples. The technique proved effective in the relative quantification of alkaloids from wild and commercial samples of *Sceletium*. However, the authors do not present any evidence validating the method in terms of linearity, limits of detection and repeatability. Nevertheless, this technique provides novel opportunities to study samples that potentially have diastereomers and isobaric structures.

**Fig. 7:**
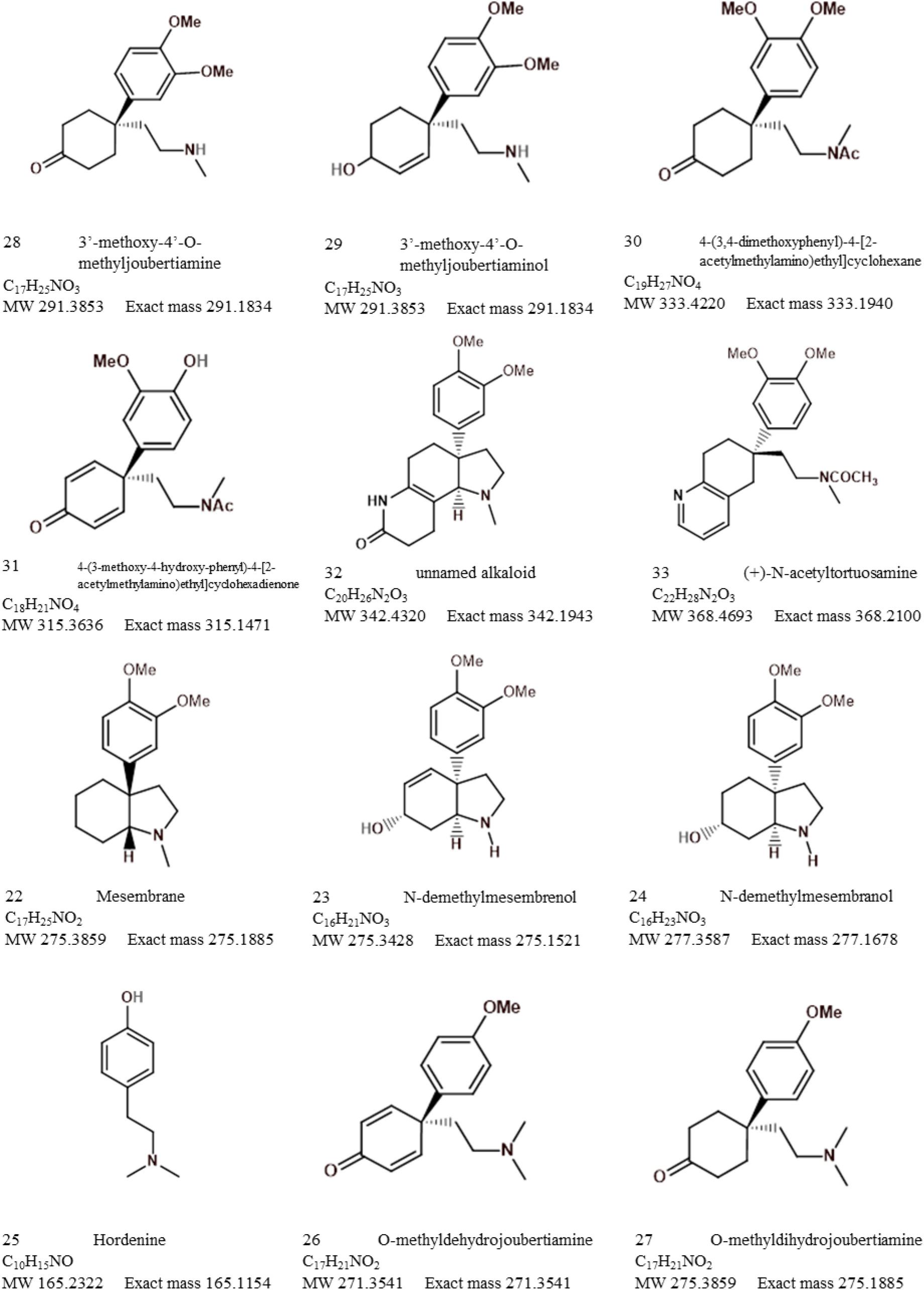
Other alkaloids found in Sceletium species (cont.)

**Fig. 8:**
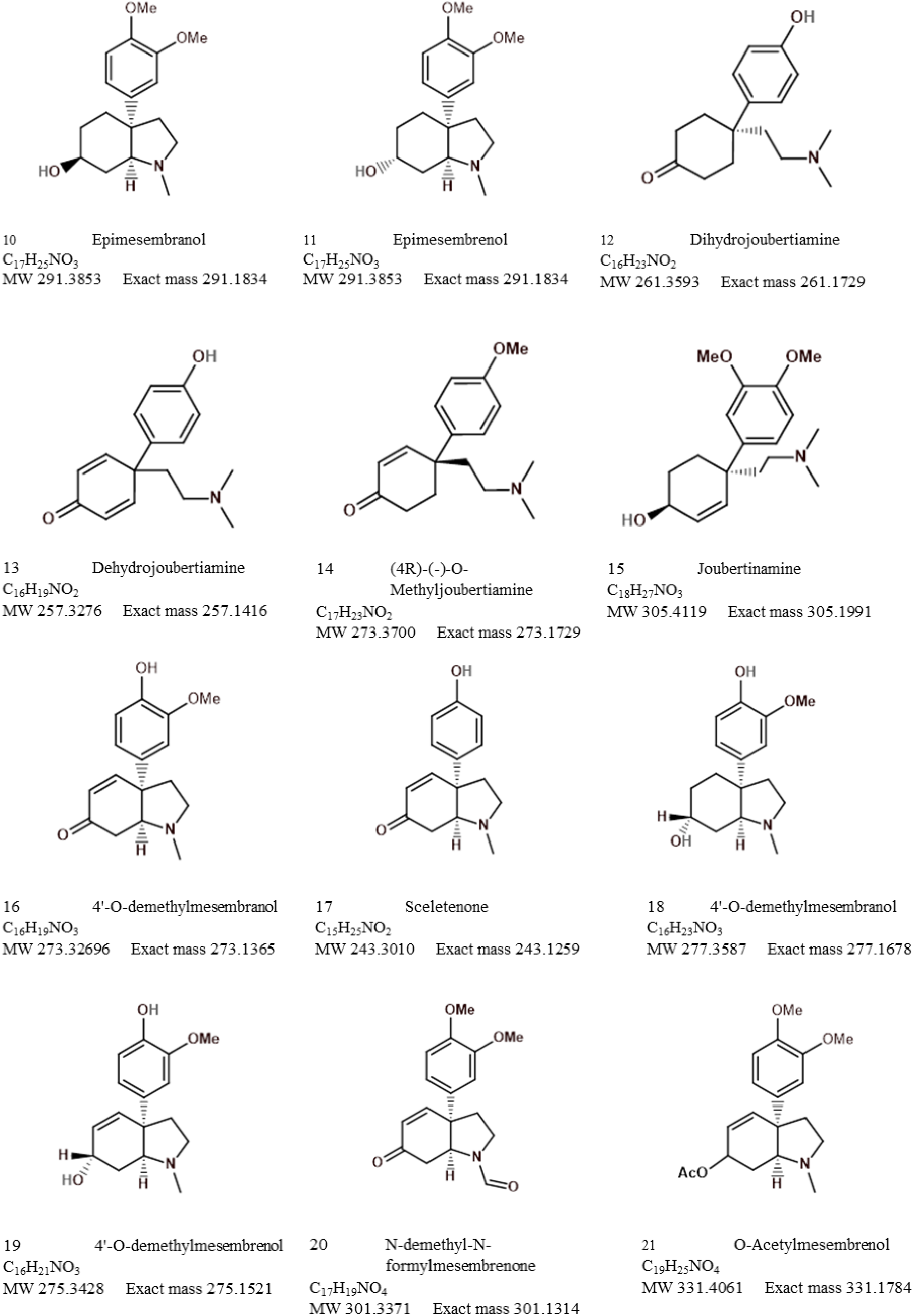
Other alkaloids found in Sceletium species

**Fig. 9:**
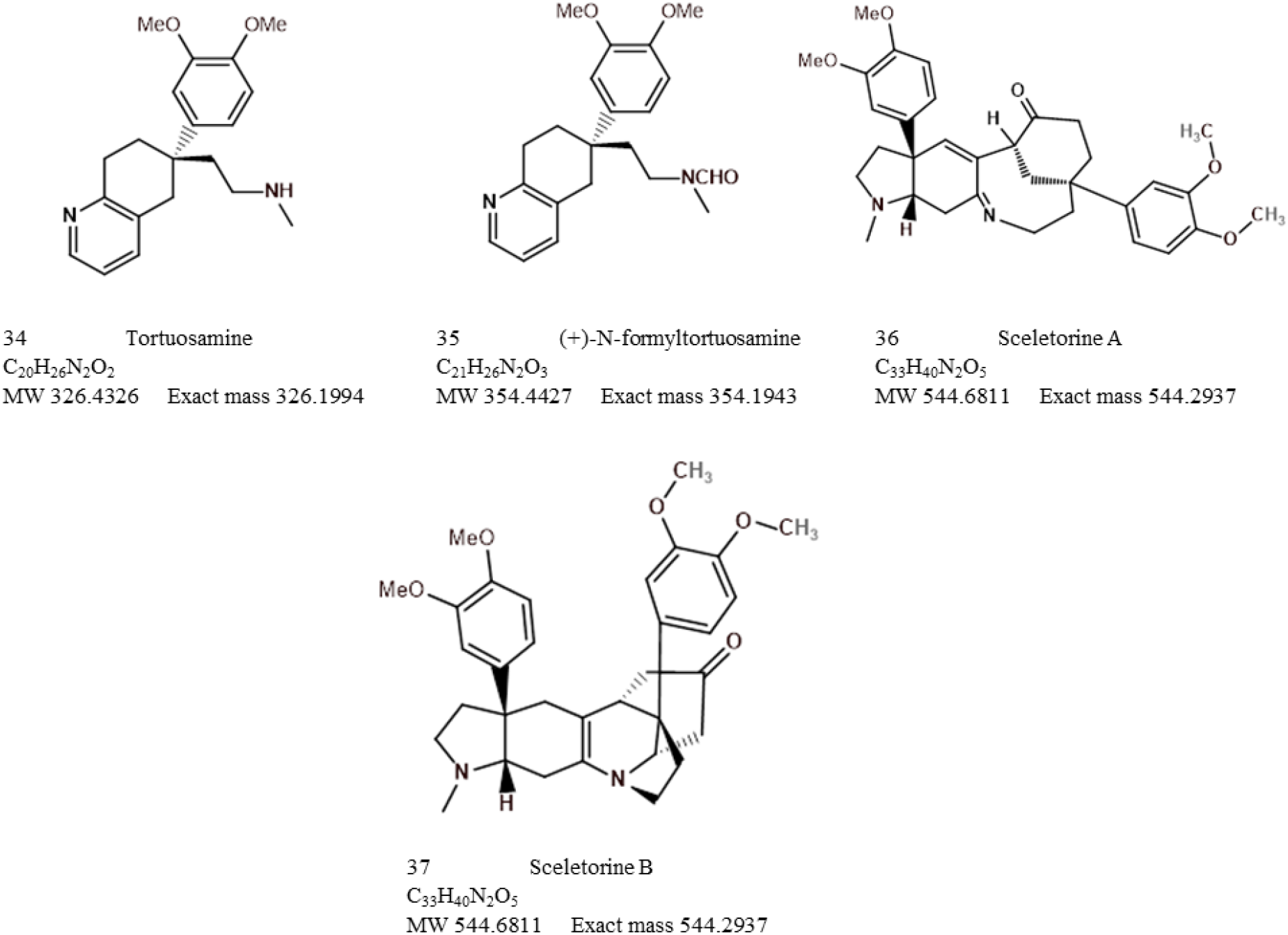
Other alkaloids found in Sceletium species (cont.)

Shikanga et al. (2013b) performed a study using UPLC and hyperspectral imaging to distinguish between *S. tortuosum* and *S. crassicuale* as these species are difficult to distinguish from each other as they look almost identical, often leading to their misidentification. Hyperspectral imaging is proving a valuable tool in the authentication of herbal products, but it is heavily reliant on good statistical models to make predictions after test materials have been scanned. Its main advantage is that it circumvents an extraction step using organic solvents, making it time efficient, non-destructive and friendly to the environment and users. This was the first study to investigate the chemical composition of *S. crassicuale.* The purpose of this study was to offer an additional robust tool to reduce the adulteration of *Sceletium* with species that may contain fewer alkaloids of interest and ultimately assist in the authentication of *Sceletium* material. The purpose of the study was to determine whether a chemometric model would offer an additional robust tool to examine unintentional adulterations of *S. tortuosum* with *S. crassicual*e. The chemical composition of *S. crassicuale* was largely uncharacterized when the authors conducted this work, and this particular species may contain fewer alkaloids of interest. The plants were dried at 30°C for two weeks after which the material was pulverised and extracted by an acid-base extraction (Shikanga et al., 2012b) (previously discussed in detail elsewhere above). The hyperspectral imaging analysis was conducted using a SisuChema shortwave infrared (SWIR) pushbroom hyperspectral spectrophotometer, with a two-dimensional array mercury–cadmium–telluride detector. The light source used was a dual quartz halogen lamp (spectrum 920–2514 nm). With an exposure of 3.0 μs, images were captured with a resolution of 10 nm. Samples were placed on a mobile plane that moved horizontally below the camera, where the full spectrum of an image was measured by line-scanning across the samples. Images were corrected for white and dark references before analysis. A multivariate principal component analysis (PCA) was created; and, background noise and edge effects were eliminated. After reference images were taken for *S. tortuosum* and *S. crassicaule*, the external samples were analysed using hyperspectral imaging. Together with this, phytochemical extracts were profiled using a previously developed method by Shikanga et al. (2012d).

The column used was a Waters Acquity reversed phase UHPLC BEH C_18_ (2.1× 150 mm, 1.7 μm particle size) column and a Van Guard pre-column (2.1× 5 mm, 1.7 μm). The injection volume was 1 μL and the temperature was 25 and 30°C, respectively. The flow rate of the mobile phase was 0.3 mL/min and the mobile phase solvent combination was (a) 0.1% aqueous ammonia and (b) acetonitrile. The UPLC-based method was thus compared to hyperspectral imagining to determine the robustness of the latter mentioned technique. The hyperspectral method illustrated that it was substantially efficient in the chemotype and chemotaxonomic classification of *S. tortuosum* and related species. The UPLC method, although widely used for the authentication of *Sceletium* samples, was not able to distinguish between *S. tortuosum* and *S. crassicuale,* possibly due to their similar chemical fingerprints as well as chemotypic variation amongst populations. However, hyperspectral imaging in combination with chemometrics proved to be an effective and robust tool in differentiating between the two species.

Patnala and Kanfer, (2013) performed a chemotypic study on mesembrine-like alkaloids that occur in the *Sceletium* genus. This study aimed to develop a technique to distinguish between samples of *Sceletium* as a quality control tool. Six species were selected based on venation patterns as distinguishing morphological characteristics are often used in taxonomy to assign species identities. Selection of plant material was based on leaf venation, with plants being grouped into the ‘tortuosum’ (*S. tortuosum*, *S. expansum* and *S. strictum*) or ‘emarcidum’ type (*S. emarcidum*, *S. exaltum* and *S. rigidum*). The species of *S. varians* and *S. archeri* were not considered in this study, in fact, these species have largely been ignored in terms of their phytochemical profiles. Plant samples were obtained from being grown in cultivation, dried at 80 °C and stored for extraction. Samples were extracted with methanol, sonicated and filtered with a final concentration of ∼150 mg/ml. The validated HPLC protocol in Patnala and Kanfer, (2010) was utilised for the quantitative determination of alkaloids of interest followed by ESI-MS and UV detection. The ‘tortuosum’ type plants, *S. tortuosum* and *S. expansum* were predominantly characterized by the presence of mesembrine, mesembrenone, mesembranol and epimesembranol. *S. strictum* was found to contain measurable amounts of mesembrine, mesembrenone and one of two epimers, 4’-O-demethyl-mesembrenone or 4’-O-demethyl-mesembrenol but mesembranol and epimesembranol were in minute relative amounts. Interestingly, the ‘émarcidum’ types illustrated a complete absence of the mesembrine class of alkaloids traditionally associated with *Sceletium* such as mesembrine, Δ^7^mesembrenone, mesembrenone and mesembranol. Instead, the émarcidum group had O-demethyl-mesembrenone and O-methyl-joubertiamine as the more prominent metabolites and out of the émarcidum’ types, *S. exaltum* showed an accumulation of mesembrine. The findings of this study clearly indicate that the distribution of mesembrine-type alkaloids is not distributed across the genus and is limited to only a few species, highlighting the importance of quality control testing in the *Sceletium* genus.

It is hypothesized that the *Sceletium* genus may have recently diversified, with minimal time between speciation events (Klak et al., 2007). As a result, these species have had a very brief period of time to accumulate differences in their DNA and subsequently are very similar in morphology. Little information is currently available with respect to the chemical fingerprints of both *S. crassicaule* and *S. emarcidum*. Patnala and Kanfer, (2015) analysed wild material of *S. crassicaule* and *S. emarcidum* using electrospray ionization mass spectrometry (EI-MS) and LCMS to characterize the chemical fingerprints of specific *Sceletium* alkaloids as a tool for the qualitative identification of lesser investigated alkaloids with complex matrices such as Δ^7^ mesembrenone, Sceletium A_4_ and epimesembranol. Furthermore, the study assessed the potential of the analytical method as a tool in quality control of *Sceletium* commercial products as tablets derived from *S. tortuosum* were included in the analysis. Their technique successfully identified Δ^7^mesembrenone, mesembrenol, mesembrenone, sceletium A_4_, mesembranol epimesembranol and mesembrine from several species of *Sceletium*.

The column used in the HPLC was a Phenomenex Luna® C_18_ HPLC column packed with 5 μm ODS-2. The elution rate was 1.0 ml/min where the elution volume was 20 μl. The detectors used were MS and UV detectors. The total run time was 16 min and the analytical procedure gave good separation of the components. The alkaloids were analysed by electrospray ionization (ESI) MS and MS/MS utilising an ionizing medium of 0.1% ammonium hydroxide in water mixed with acetonitrile at a flow rate of 0.3 ml/min. The conditions of the ESI-MS system were: a capillary temperature of 240 °C and a spray voltage maintained at 4.5 kV for all compounds. Within the MS/MS mode, collision induced dissociation (CID) was induced for the detection of protonated molecular ions [M+H]^+^, enabling a better collection of fingerprints for each alkaloid in this system. The method identified Sceletium A_4_ from *S*. *crassicaule* material. Of interest, *S. emarcidum* did not have any of the reference compounds, that normally occur in *Sceletium* samples. The investigation did not report on any dominant structures that could be used for the chemotaxonomic classification of this species, as none of the peaks observed corresponded with the standard alkaloids found in *Sceletium* species. The versatility of this method may be of value in characterising other alkaloids from *Sceletium* aside from the mesembrine-type alkaloids. It has the power to be employed routinely for a chemotaxonomic delineation of *Sceletium* species because it can separate different isomeric derivatives that are often neglected or difficult to define in the chemical analysis of *Sceletium* phytoextracts. Unlike the UPLC-UV method (Patnala and Kanfer, 2010) and the RP-UHPLC method (Shikanga et al., 2012b), the ESI-MS method was able to identify Δ^7^mesembrenone and epimesembranol. It was effective in qualitatively differentiating isobaric, isomeric and epimer compounds. Alkaloids of this nature often prove a challenge due to their similar chemical structures, where mesembrenone and Δ^7^mesembrenone differ by the position of a double bond (isometric) and mesembranol and epimesembranol differ by configurations linked to epimer or diastereomer formation.

Apart from plant misidentification and the choice of inferior chemotypes that express poor bioactivity, chemical and heavy metal adulterations as well as herbal adulterations of phytomedicines can lead to undesired cytotoxic effects upon human consumption. Metabolite profiling can thus be a complimentary tool to other techniques for the detection of adulterants in herbal medicines. To this end, Lesiak et al. (2016) performed analysis on *S. tortuosum* commercial material using direct analysis in real time ionization coupled with high resolution time-of-flight mass spectrometry (DART-HRTOF-MS) in positive ion mode. The method was employed as an authentication tool to identify adulterated samples and found that some commercially available samples were indeed spiked with the banned herbal stimulant ephedrine. The settings of the DART ion source were set with a heater temperature of 350 °C, grid voltage of 250 V and helium flowrate of 2.0 L.s^-1^. Commercial powder mixtures were conveniently analysed directly by dipping the closed end of a melting point capillary tube into the powder substance and then between the DART ion source and mass spectrometer inlet. The analytical technique was able to identify the following alkaloids: mesembranol, mesembrine, dihydrojoubertiamine, O-methyljoubertiamine, O-methyldehydrojoubertiamine, sceletenone, dehydrojoubertiamine, 4-O-desmethylmesembrenone, 4-O-desmethylmesembrenol, 4-O-desmethylmesembranol and hordenine. The authors only quantified two of the detectable compounds, mesembranol ranging from 0.3 to 7.0% and mesembrine at 5.1% but relative amounts are not available for any of the other compounds in the authors’ report. Quantification of *Sceletium* alkaloids is often constrained by the availability of pure reference standards. Despite this, this analytical tool displayed sufficient capability of searching for other alkaloid types within the *Sceletium* samples. It also provided a rapid forensic diagnostic tool of commercial samples sold in the USA, highlighting illicit practices in the manufacture of *Sceletium*-derived products that are of regulatory concern.

Appley et al., (2022), was another analytical study concerned with the authentication of *Sceletium*-based products through scientific verification using the biomarker hordenine and several mesembrine alkaloids to develop a reproducible protocol for the forensic analysis of products containing *Sceletium*. Supporting the method used by Lesiak et al. (2016), the use of direct analysis in real time–high-resolution mass spectrometry (DART-HRMS) in positive ion mode, resulted in effective and rapid detection and quantification of hordenine and mesmebrine-type alkaloids. The settings of the DART ion source were set with a heater temperature of 350 °C, grid voltage of 250 V and helium flowrate of 2.0 L.s^-1^. Extracts were prepared from commercial *Sceletium-*based products. The analytical method was able to resolve hordenine as a biomarker in all commercial samples along with the mesembrine-type alkaloids, mesembrenone, mesembrine, mesembrenol and mesembrinol. The study provided a reproducible and robust protocol for the quantification of hordenine and mesembrine-type alkaloids from *Sceletium* that could support the forensic relevance of the natural product’s use.

The main concern with the presence of ephedrine in natural products is that it has been noted as being fatal when combined with caffeine or other over-the-counter drugs and could potentially be harmful when consumed with *Sceletium* products (Haller and Benowitz, 2000). An advantage of DART-HRTOF-MS is that sample preparation is not needed. As a consequence, there is no loss of phytochemical constituents due to solvent bias as samples are analysed in their unaltered form. Techniques such as LC-MS and GC-MS may not identify adulterants such as ephedrine due to preferential take up of hordenine due to its polarity over the adulterant ephedrine as these are constitutional isomers of each other, which both occurred at a nominal *m/z* of 166. The authors emphasised that without the use of DART-HRTOF-MS, the compounds would not have been separated and the adulterant would have thus become more difficult to notice and identify. This study illustrated the utility of the analytical tool for rapid identification of adulterants, specifically ephedrine, in *Sceletium*.

The advantages of using NMR for generating an overview of the plant metabolome lie in its vast applications ranging from quality control of foods and botanicals to studies related to investigating the pharmacological activity of phytochemicals. Furthermore, the use of NMR spectroscopy is appropriate for samples where authentic standards are not available (Leiss et al., 2011) for a wide range of metabolites that may occur within a particular plant specimen, such as the case with *Sceletium*. Despite this NMR spectroscopy does have limitations in some cases as without the use of two-dimensional NMR, absolute quantitation is thus not possible (Verpoorte et al., 2007). Although not too often, the use of NMR for metabolomic analysis of *Sceletium* plants is indicated in the primary literature.

Zhao et al. (2018) used both ^1^H-NMR spectroscopy and ultra-performance liquid chromatography-mass spectrometry (UPLC-MS) to analyse wild samples of *S. tortuosum* in an effort to elucidate chemotypic variation between two populations groups (Western Cape and Northern Cape) over 23 localities in South Africa, using a metabolomics-driven strategy (Table 1). The populations were collected from various locations over the Western and Northern Cape. The study of plant metabolomes has seen an unprecedented rise since the adoption of systems biology approaches in biological sciences and NMR-metabolomics can be the preferred choice for this purpose (Verpoorte et al., 2007; Leiss et al., 2011). The advantage of using NMR for generating an overview of the plant metabolome lies in its vast applications, ranging from quality control of foods and botanicals to studies related to investigating the pharmacological activity of phytochemicals. A limitation of NMR spectroscopy is that without the use of two-dimensional NMR, absolute quantitation is not possible (Verpoorte et al., 2007). The plants collected at different localities studied by Zhao et al. (2018) using NMR were growing under differing biogeographic environments. The Western Cape samples were collected from semi-desert, dry zones with winter to spring rainfall. On the other hand, the Northern Cape samples were collected from a similar climatic region that is also known to be semi-arid with summer rainfall. There were also other samples collected from moist areas that would have experienced rain in both winter and summer. For NMR, plant samples (200 mg each) were extracted with 500 mL deuterated methanol (CD3OD, 99.8% D). The NMR instrument was run at 499.79 MHz for ^1^ H and 125.67 MHz for ^13^C NMR. Samples were sonicated and centrifuged. Methanol was also used for extraction purposes of the harvested plants before GC-MS. Gas chromatography settings were performed on a fused silica capillary column (30 m 0.25 mm i. d x 0.25 mm film thickness) with a carrier gas of helium (1.0 mL/min). For the UPLC method, the extraction method used in this study was the same as that of Shikanga et al. (2012d). The column used in the unit was a C_18_ column (150 mm 2.1 mm, i.d., 1.7 mm particle size) at a temperature of 30 °C. The mobile phase was 0.1% ammonium hydroxide (Solvent A) and 90% acetonitrile (solvent B) at a flow rate of 0.3 mL/min. The study found that NMR together with a chemometric analysis could be an effective tool to distinguish between populations of *Sceletium* and identify notable biomarkers in each population. A significant finding from the study was that *N*-demethyl-*N*-formylmesembrenone, a biomarker that had not been identified in *Sceletium* before, characterised one of the population groups from the Western Cape. Separation into groups was achieved by differences identified in biomarker groups of pinitol, alkaloids and alkyl amines. The study reported that the production of alkaloids may be due to genetic composition rather than climatic conditions since plants in close proximity to each other produced variable amounts of alkaloids, suggesting that climate was not a contributing factor to diversity in chemical profiles. The findings from this study suggested that the collection of *Sceletium* for commercial purposes should be done from cultivated samples instead of wild samples, due to the highly variable alkaloid distribution across populations that was noticeable between and within groups. This alkaloid variability was predominantly observed in the Western Cape samples where 2.8% of samples were devoid of any of the four mesembrine alkaloids of interest, namely, mesembrine, mesembrenol, mesembrenone and mesembranol. These data correlated with reports from Shikanga et al. (2012d) where 4.2% of samples from the Western Cape were lacking in mesembrine alkaloids. The Northern Cape populations were more consistent in their chemical profiles between studied samples.

Freund et al. (2018) employed an analytical technique that had never been performed on these plants coined leaf spray mass spectrometry (leaf spray MS) (Table 1). Additionally, the setup included tandem mass spectra (MS/MS) collected in positive ionization mode. This technique circumvents the separation of plant metabolites using chromatography and offers a direct MS injection without the need for sample preparation or extraction, with minimal technical adjustments to the ionization source being required. An advantage of this tool is the absence of lengthy extraction protocols and solvent bias that may introduce artifacts or prove inefficient in the recovery of phytochemicals as these may not always be extracted. Leaf spray MS can directly analyse plant tissue providing rapid generation of qualitative and quantitative data albeit accurate quantification is more tricky with this approach. The setting for the MS system was: sheath, auxiliary, and sweep gas to 0; the spray voltage to 2 - 5 kV; the capillary temperature to 150 - 250 °C; and the S-lens RF level to 50. The phytochemicals putatively identified from *S. tortuosum* material were, dihydrojoubertiamine (m/z 262.1794), O-demethyl-mesembrenone (*m/z* 274.1431), O-demethyl-mesembrine (*m/z* 276.1583), N-demethyl-mesembrine (*m/z* 276.1583), O-demethyl-dihydro-mesembrenone (*m/z* 3 276.1583), N-demethyl-dihydro-mesembrenone (*m/z* 276.1583), mesembrenone (*m/z* 288.1587), mesembrine (*m/z* 290.1742) and dihydro-mesembrine (*m/z* 292.1897). The method was successful in analysing intact plant material reducing the amount of processing needed. For leaf spray MS to have wider application for *in planta* analysis of metabolites, optimisations in terms of plant preparation, presence or absence of solvents, volume of solvents, voltage amplitude and distance from the ion inlet may be necessary. Some other limitation of this analytical technique is its low dynamic range, resulting in the most abundant metabolites solely being identifiable. Some of the minor alkaloids or structurally similar alkaloids that require stronger resolving power require greater and more sophisticated technical expertise for their detection. Despite this, the analytical method proved to be a powerful tool that eliminated chromatography for the identification of the main phytochemicals from *S. tortuosum*.

Sandasi et al. (2018) performed a study to assess the quality of herbal tea blends using hyperspectral imaging and UPLC-MS, in an effort to develop a tool for the quality control of various *Sceletium* tea blends. Five batches of herbal tea that were claimed by the manufacturers to contain a *S. tortuosum* and *Cyclopia genistoides* (commonly known as honeybush) mixture were obtained pulverised and subjected to hyperspectral imaging without any further processing. For UPLC-MS analysis, the tea blends were prepared by adding boiling water (237 ml) to 1.5 g of plant tissue. The test specimens were stirred continuously for 25 minutes before 2 μl was injected into the UPLC instrument. The column used was C_18_ column (150 mm × 2.1 mm, i.d., 1.7 μm particle size) maintained at 30 °C. The solvent system was Solvent A: 0.1% ammonium hydroxide Solvent A: 0.1% formic acid Solvent B: 90% acetonitrile at a flow rate of 0.3 ml/min. A gradient elution was applied with a total run time of 8 minutes. The electrospray mode was set to positive ion. For the hyperspectral imaging, the same instrument settings were used as in Shikanga et al. (2013b) and applied as a rapid and non-destructive method for the quality control of the tea blends. Using a PLS-DA model, the procedure had a 95.8% predictive ability providing high degrees of sensitivity with a stronger metabolite feature selection. The five tea blends were predicted to only contain *S. tortuosum* and *C. genistoides* with no adulterants or contaminants. Quantitatively, *C. genistoides* was found to be in higher amounts across the samples (> 97%), while *S. tortuosum* was found in lower quantities (< 3%). After UPLC-MS, the chemometric technique confirmed the identity of *S. tortuosum* and facilitated the identification of three out of the four alkaloids that occur in this species that were of interest. The alkaloids found were: mesembrine (*m/z* 290.1609), mesembrenone (*m/z* 288.1460) and mesembranol (*m/z* 292.1764). The absence of the fourth commonly occurring alkaloid may be due to it eluting with other peaks becoming obscured during analysis but the role of chemotype variability in the samples could not be ruled out (Roscher et al., 2012; Shikanga et al., 2013b). For this study, the UPLC-MS conditions were optimised for *C. genistoides* but these conditions were not necessarily optimal for *S. tortuosum*. A major limitation of this particular study was that the UPLC-MS procedure alone could thus not conclusively efficiently distinguish the plant components of the herbal mixtures that contained *S. tortuosum* along with another species. Additionally, the quantitative determination of phytochemicals in a mixture may be biased depending on which protocol of analysis is being employed. Combining hyperspectral imagining with chemometrics proved a more powerful and reproducible tool for the quality control of herbal tea blends containing *Sceletium* and honeybush.

It is not clear why there are so few reports on the isolation and characterisation of channaine, however, it is likely that analytical methods being used by researchers are not necessarily optimised for the detection of this unusual alkaloid channaine. We speculate that this compound may also be produced at minor levels during the life time of the plant, making it even more difficult to isolate. Veale et al. (2018), aimed to structurally elucidate the alkaloid, channaine from *S. tortuosum*. This was interestingly the second time that channaine was detected since its initial characterisation by Abou-Donia et al. (1978). The plant extract was prepared by collecting macerating the aerial parts of *S. tortuosum* and air drying the material prior to an acid-base extraction Shikanga et al. (2011). Isolation of channaine was achieved through preparative UPLC in tandem with a quadrupole time-of-flight (Waters Xevo® G2QToF) mass spectrometer. The detector used was a PDA detector. The column used was an XBridge Prep C_18_ column (19 × 250 mm, i.d., 5 μm particle size) maintained at 40 °C. The injection volume was 100 μL and mobile phase 0.1% ammonium hydroxide in water (Solvent A) and acetonitrile (Solvent B) at a flow rate of 20 mL/min. Fractions were collected and analysed on a 600 MHz NMR apparatus. This was the first full NMR analysis of channaine in the literature. Chemical structures were resolved using ^1^ H, ^13^C, COSY, HSQC and HMBC NMR spectroscopy.

The search for novel chemicals from *Sceletium* species has found renewed interest. Recently, Yin et al. (2019) performed an extraction of *S. tortuosum* and isolated sceletorines A and B for the first time with these authors making suggestions on plausible biosynthetic pathways associated with sceletorine production. This kind of information is largely missing in terms of novel alkaloids that become periodically identified in *Sceletium* samples by different research groups. The isolated alkaloids were established to be precursors of the alkaloid channaine identified in previous studies (Abou-Donia et al., 1978; Veale et al., 2018). It was ruled out that these phytochemicals were artefacts as a result of processing due to their presence in fresh material. Aerial parts of *S. tortuosum* (1.5 kg) were extracted with methanol (4 L×24 h × 4). An acid-base extraction was then performed; 5% HCl in water (1.3 L) followed by EtOAc (1.3 L × 3). Preparative TLC (20 cm × 20 cm, 500 μm) and column chromatography (silica) were conducted. The solvent system used in the column was CHCl_3_−CH_3_OH–NH_4_OH (from 100:0:0 to 0:100:0.3). Sub-fractions were obtained by using solvent systems of CHCl_3_, CHCl_3_−CH_3_OH–NH_4_OH (300:7.5:1) and (2000:100:7). Alkaloid fractions were visualised with Dragendorff’s and vanillin visualisation stains. Semi-preparative HPLC analysis was performed with a UV-Diode detector. The column used was a RP-C_18_ column (250 × 10 mm; particle size 10 μm). Acetonitrile and water were used as the solvent system (A (water with 0.1% acetic acid) and B (acetonitrile with 0.1% acetic acid)). The injection volume was 100 μL and a flow rate of 4.0 mL/min. Sensitivity, reproducibility and comparison to a reference method of these two new alkaloids were not presented in the paper. The authors proceed to successfully characterized in great detail by using NMR.

The main challenges associated with the purification and identification of alkaloids from *Sceletium* are associated with irreversible adsorption to column packing materials, excessive tailing, and poor recovery as well as catalytic changes encountered with solid supports in various analytical systems (Yang and Ito, 2005). These challenges have been reported to be overcome to some extent by high-speed counter-current chromatography (HSCCC) (Shikanga et al., 2011) and non-aqueous capillary electrophoresis coupled to mass spectrometry (NACE-MS) (Roscher et al., 2012). The technique of HSCCC allows for efficient separation as it is an all-liquid method allowing great recovery (Shikanga et al., 2011) while NACE-MS allowed for the sensitive identification of diastereomers in material (Roscher et al., 2012). It should be noted that the technique has not been rigorously tested for reproducibility in larger data sets. In terms of analytical techniques for rapid quality control, tools such as DART-HR-TOF-MS (Lesiak et al., 2016) and leaf spray MS (with MS/MS) (Freund et al., 2018). These techniques are robust, rapid and require very little plant material. However, it should be noted that the limitations of the latter technique are that it still requires optimization. For leaf spray MS, the succulence of *Sceletium* spp. is ideal and advantageous as its application is well suited to analyse fresh plant material containing high amounts of water, which may be a challenge for other methods.

Reddy et al. (2022), investigate the chemotypic variation across populations of *Sceletium* species. This is one of the few studies looking at the chemical composition of other species in the genus and the only one to report on the chemical composition of *S. rigidum* and *S. emarcidum*. The analytical technique of HPLC-MS-MS was employed and data were processed using Feature-based Molecular Networking to annotate the chemical space and investigate the chemical space in greater detail to identify minor and co-eluting phytochemicals. The study put forward *in silico* results supporting that minor phytochemicals identified in *Sceletium* species may be responsible for the therapeutic activities observed in the literature (Harvey et al., 2011; Krstenansky, 2017).

It is of great importance that investigators make note of the great chemotypic variation present in populations of *Sceletium* (Shikanga et al., 2012d; Zhao et al., 2018). Overall analytical techniques should be performed to assess sensitivity, reproducibility and comparison to a reference method (i.e. GC-MS). From the current state of analytical techniques used in the quality control of *Sceletium*, GC-MS, LC-MS and HPLC-MS will continue to remain popular going forward (Table 1). However, for the effective identification of adulterants and contamination in samples, more advanced tools in tandem with different detectors need to be utilized. Two analytical techniques that stand out for the rapid analysis of samples are direct analysis in real time ionization coupled with high resolution time-of-flight mass spectrometry (DART-HR-TOF-MS) (Lesiak et al., 2016) and leaf spray MS (Freund et al., 2018). These methods do not require the processing of material and as such there is no solvent bias, loss of phytochemicals during extraction, or artifacts from extraction procedures. Nevertheless, the limitations of these methods are that the machines are not common, expensive and require specialized components that may prove to be more laborious to assemble. With these in mind, the application of NMR analysis coupled with chemometrics (Zhao et al., 2018) will also gain more popularity in quality control assurance practices. Methods that are non-destructive to the plant tissues such as hyperspectral imaging may provide additional analytical power for use in commercial settings (Sandasi et al., 2016, 2018). Further advancements in analytical techniques will likely result in novel methods that may be used in the future.

### Fermentation of *Sceletium*

The fermentation of *Sceletium* and the effect on the medicinally important alkaloids has been of interest due to the traditional preparation and the anecdotal reports of the plant becoming more euphorically potent when fermented (Smith et al., 1996). The plant material is fermented for several days before being dried for storage and use (Gericke and Viljoen, 2008). Traditionally, the Khoe-Sān people of South Africa prepared the plant material by crushing and fermenting the aerial parts of *Sceletium*, which was then placed in an animal skin bag for several days (Chen and Viljoen, 2019). It has been reported that in recent years, plastic bags are used as an alternative to canvas or skin bags (Van Wyk and Gericke, 2000). The mood-elevating activity is currently suspected to be linked to four mesembrine-type major alkaloids, mesembrine, mesembranol, mesembrenone, and mesembrenol (Gericke and Viljoen, 2008). Fermentation is thought to enhance the levels of these alkaloids and reduce oxalates which in turn increases the mood-elevating activity of *Sceletium* (Smith et al., 1996). There seems to be some incongruency in reports related to the effects of fermentation which further highlights work that needs to be done to better understand the metabolic pathways of these alkaloids and at which steps to manipulate levels.

Many analyses have investigated the effect of fermentation on the alkaloid profile in *S. tortuosum* (Table 2). Smith et al. (1998) used the plastic bag fermentation followed by GC-MS analysis. Their findings indicated that the mesembrine alkaloid composition was comparable to that of the oven-dried (80 °C) samples. However, in the fermented sample, there was a significant increase in mesembrenone levels whilst levels of 4’-*O*-demethylmesembrenol and mesembrine decreased. Patnala and Kanfer, (2009) further investigated the role of fermentation in alkaloid composition in *Sceletium* using HPLC-MS, where levels of mesembrine decreased (from 1.33 to 0.05% (w/w)), with suspected transformation into Δ^7^mesembrenone. Roscher et al. (2012), also investigated the change in alkaloid composition as a result of sample fermentation. No qualitative data were presented in this study instead the authors report that no overall change in the alkaloid concentrations as a result of fermentation were detected. The studies conducted up until this point were suggestive that the traditional processing of the plant material through fermentation did not affect the overall potency of the material as there was a decrease or no change in mesembrine levels (Smith et al., 1998; Patnala and Kanfer, 2009; Roscher et al., 2012). However, the most recent study by Chen and Viljoen, (2019), reported that the total alkaloid content increased as a result of fermentation. They reported that while mesembrine levels increased (from below 1.6 μg/mL to 7.40–20.8 μg/mL), there was only a marginal increase in mesembrenol and mesembranol content and a significant decrease in mesembrenone content. This study supports the traditional preparation of *Sceletium* plant material to increase the mood-elevating effects of *S. tortuosum.* It may be worthwhile to investigate if phytochemical formation or breakdown is dependent on pH in these analyses and that future analyses should control for this. Previous studies have not indicated the pH of their extracts or used them in their analyses.

**Table 2:** Methods used for fermentation and results obtained from previous research (↑ =increase, ↓ =decrease).

| Extraction method | Fermentation procedure | Analytical technique | Conclusion | Reference |
| --- | --- | --- | --- | --- |
| Classical acid-base extraction | Plants crushed with soil, placed in plastic bag and sealed in airtight plastic bag. | GC-MS | ↓ 4'-O-demethylmesembrenol<br>↓ mesembrine<br>↑ mesembrenone | (Smith et al., 1998) |
| Methanol | Aerial plant parts crushed and placed in plastic bag exposed to sun for 10 days. | HPLC-MS (validated) | ↓ Mesembrine<br>↑ $\Delta^7$- mesembrenone | (Patnala and Kanfer, 2009) |
| Methanol | Plant material stored in airtight plastic container for 8 days. | NACE-MS | Alkaloid profile remained unchanged, no quantitative data reported. | (Roscher et al., 2012) |
| Methanol | Aerial plant parts gently bruised and placed in an incubator (40 °C) for 7 days | UPLC-MS (validated) | ↑ Mesembrine<br>↓ Mesembrenone | (Chen and Viljoen, 2019) |

Comparing the fermentation studies available in the literature thus far, it is clear that there is no conclusive evidence that fermentation results in a consistent change in the alkaloid profile. More investigation is needed to understand how fermentation affects alkaloids in *Sceletium* and what the best method of fermentation is, that most accurately represents traditional ethnobotanical preparation. A better understanding of the biosynthetic pathway(s) could assist in understanding how fermentation influences the metabolite profile. Currently, the biosynthetic pathway of mesembrine-type alkaloids as suggested by Jeffs et al. (1971a), proposed that the perhydroindole portion of mesembrine comes from tyrosine and the aromatic group is derived from phenylalanine. The cinnamic acid derivatives are produced from phenylalanine but the 3’-aryl oxygen substituent is proposed to be introduced in later steps involving a biosynthetic reaction with sceletenone, mesembrenone and 4’O-demethylmesembrenone (Jeffs et al., 1978). Since studies in 1971 and 1978, the biosynthetic pathway has not been revised and we thus suggest this become a future avenue of investigation to better understand how fermentation affects the alkaloid profile, pinpointing not only biochemical changes that occur but also the key enzymes and genetic regulatory steps that may control this metabolism.

### Present-day ethnobotanical use

Although there is limited current ethnobotanical information on the prevalence of use of the *Sceletium* genus in modern times, the work by Philander, (2011) focusing on a group of Rastafarian herbalists who are the main commodifiers of traditional medicines in the Western Cape of South Africa, clearly points out to the importance of *S. tortuosum* as the main species that is collected as a phytomedicine to reduce depression and anxiety, where the plant is mainly administered through drinking it as a herbal tea or by chewing. Anecdotal evidence for its use in indigenous medicine does however exist. For example, the community of Rastafarian herbalists that belong to the Cape Bush Doctors organisation use it together with *Cannabis sativa*, for spiritual purposes (Olivier, L, personal communication). More recently, in popular culture, *Sceletium* has been used for recreational for its mood-elevating and anxiolytic effects. With the increased public interest in biogenic drugs such as *Sceletium*, numerous companies have appeared online selling *Sceletium* in raw powdered form, tablets, teas and snuffs. This increased popularity may well pose a significant conservational threat to the species if populations are collected from the wild. As evident in multiple studies, distinguishing between the morphological characteristics can be challenging (Patnala and Kanfer, 2010) without the consultation of specialised botanists or analytical tools. It appears that some of the present-day uses by indigenous communities of the plant are consistent with the historical uses i.e. for euphoria and as a mood elevator (Smith et al., 1996; Gericke and Viljoen, 2008). However, it is unclear if more rural communities where these plants are found are still relying on using these plants for hunger and as a thirst suppressant.

### Biological activities of *Sceletium* extracts and isolated alkaloids

There are many claims in the literature that report the use of *Sceletium* to suppress hunger and thirst but at this particular stage, there are limited scientific reports that focus on animal models to test such claims are available. There are, however, several reports on the ‘mood elevation’ activity, particularly focussed on the potential of *Sceletium* to aid with anxiety and depression. Recently, there have been more studies that are focused on the pharmacology of *Sceletium*, notably in areas linked to *in vivo* actions and clinical trials of tested extracts (Table 3). The body of pharmacology research on *Sceletium* is quite extensive, as seen below, with the majority of the reported biological activity being interactions with the central nervous system (CNS) and related neurological pathways (anti-depressant, anxiolytic and psychoactive activity). The scope of the observed CNS-activity is broad with observed anxiolytic and anti-depressant activity demonstrated for extracts and isolated compounds of *Sceletium* (Table 3). Additionally, there are more *in vivo* behavioural inquiries on rats using a range of pharmaceutical tests (Fig. 5d) exhibiting CNS-related activities ranging from suppressant (e.g. anxiolytic and sedative) to excitatory (e.g. antidepressant) activity. Although there is a shift to more *in vivo* studies, there is still much to be tested in terms of different chemotypes and to understand the pharmacokinetics of individual phytochemicals and potential synergism between phytochemicals, aside from mesembrine.

**Table 3:**
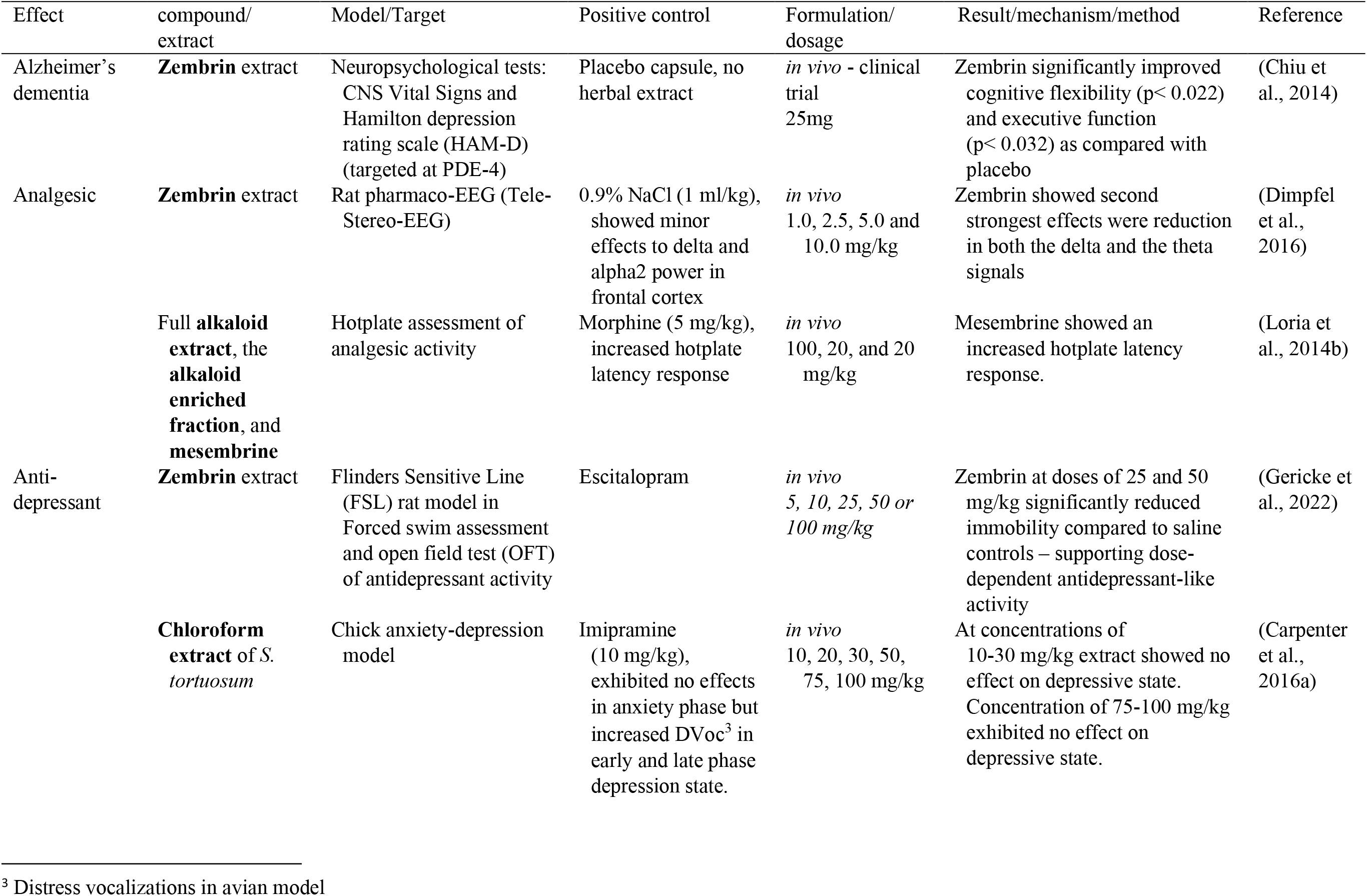
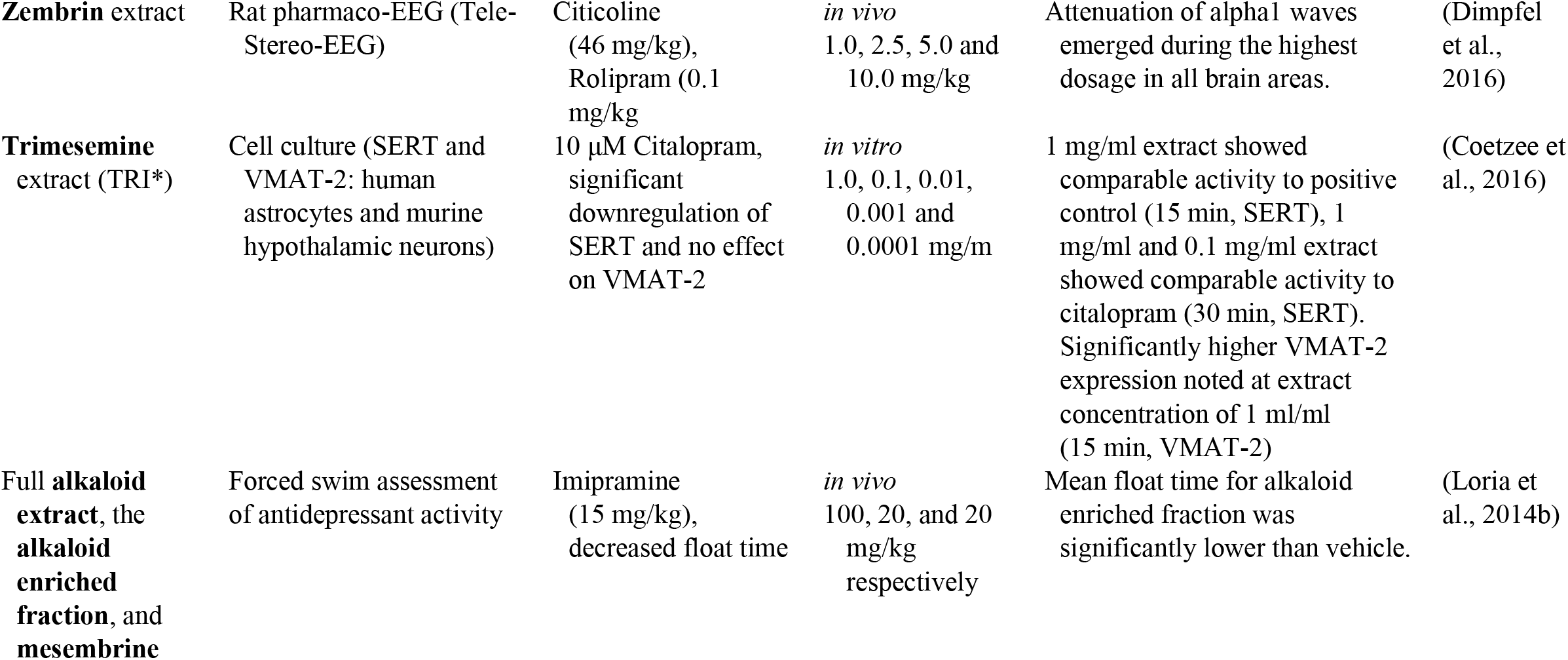
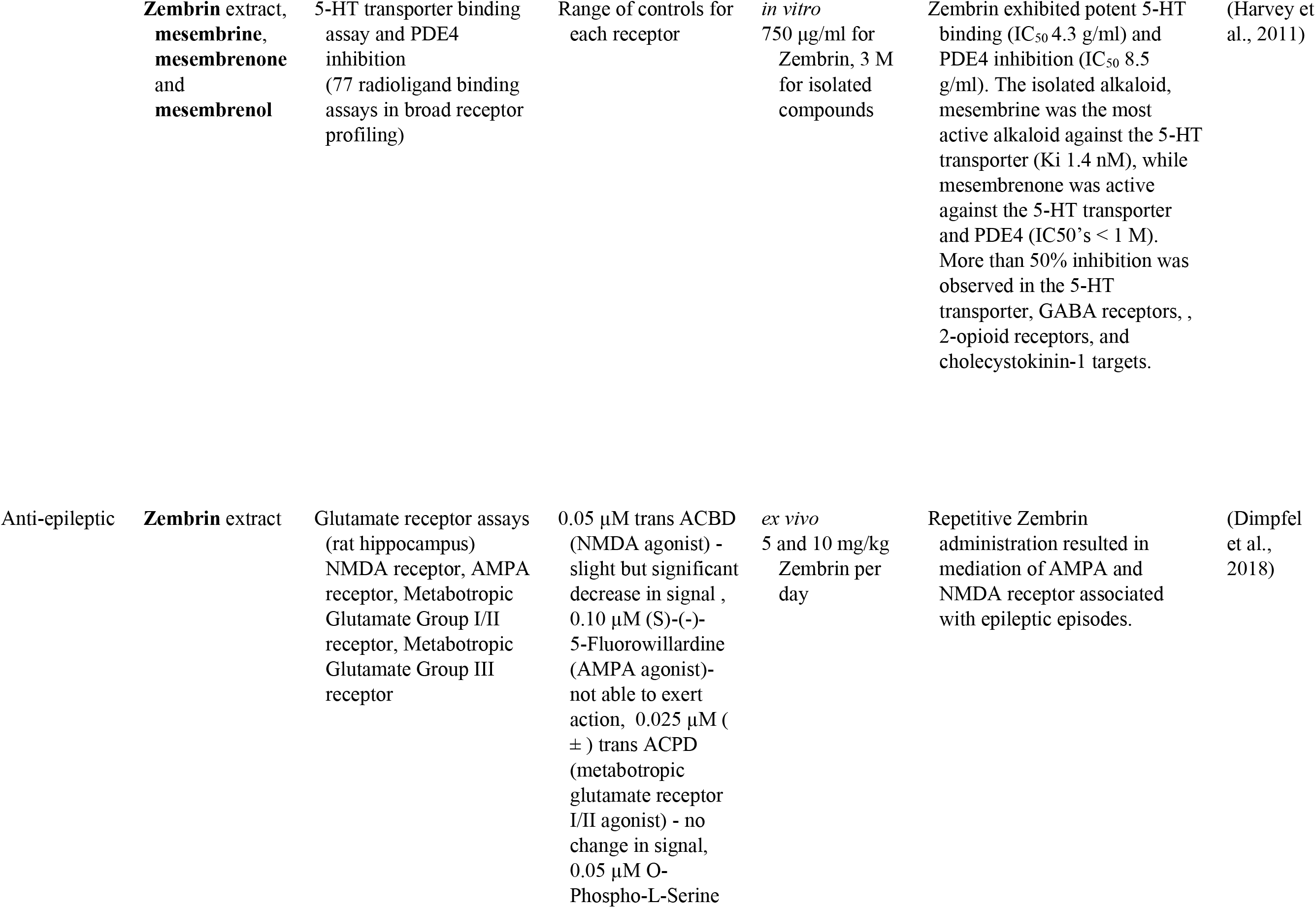
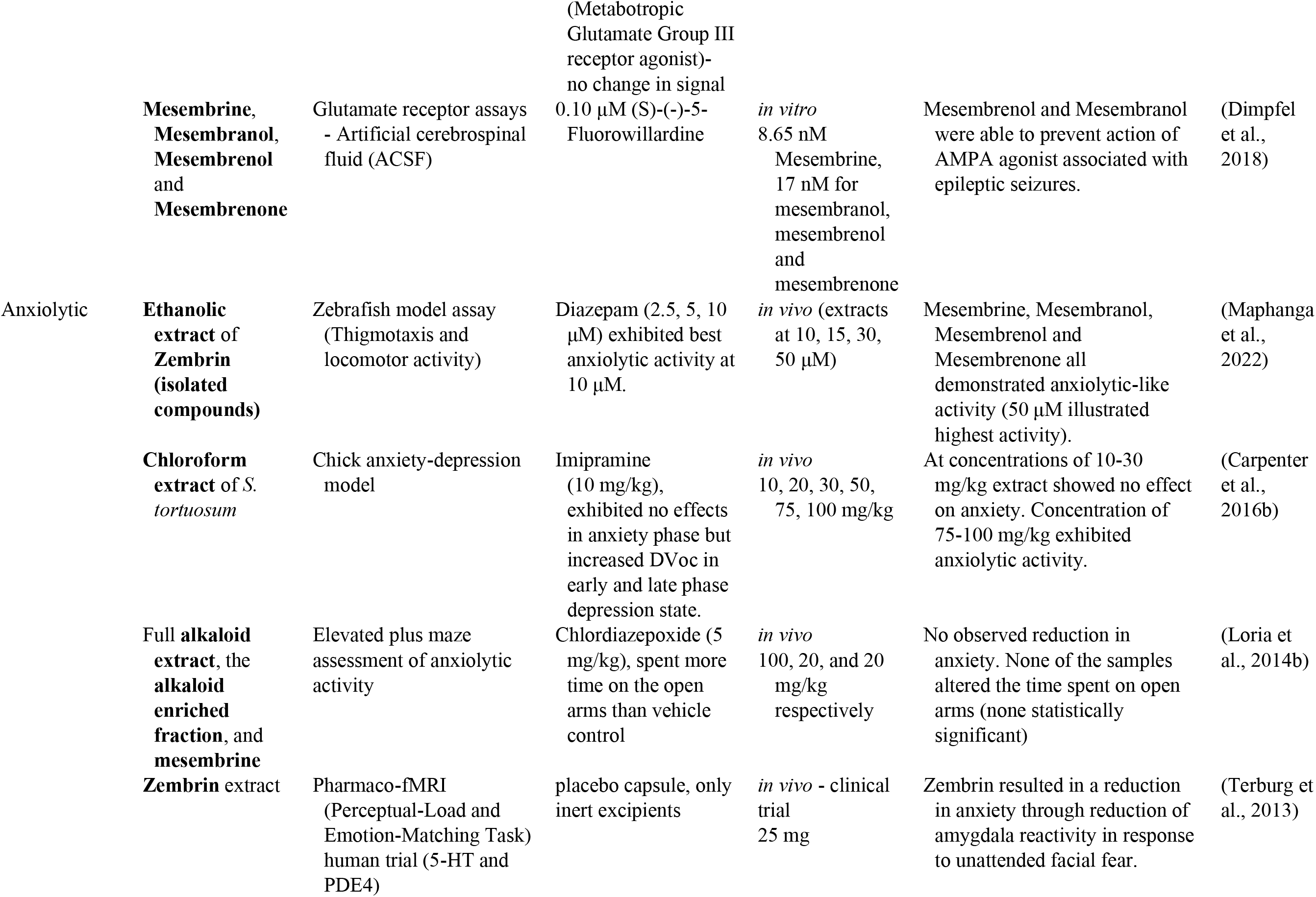
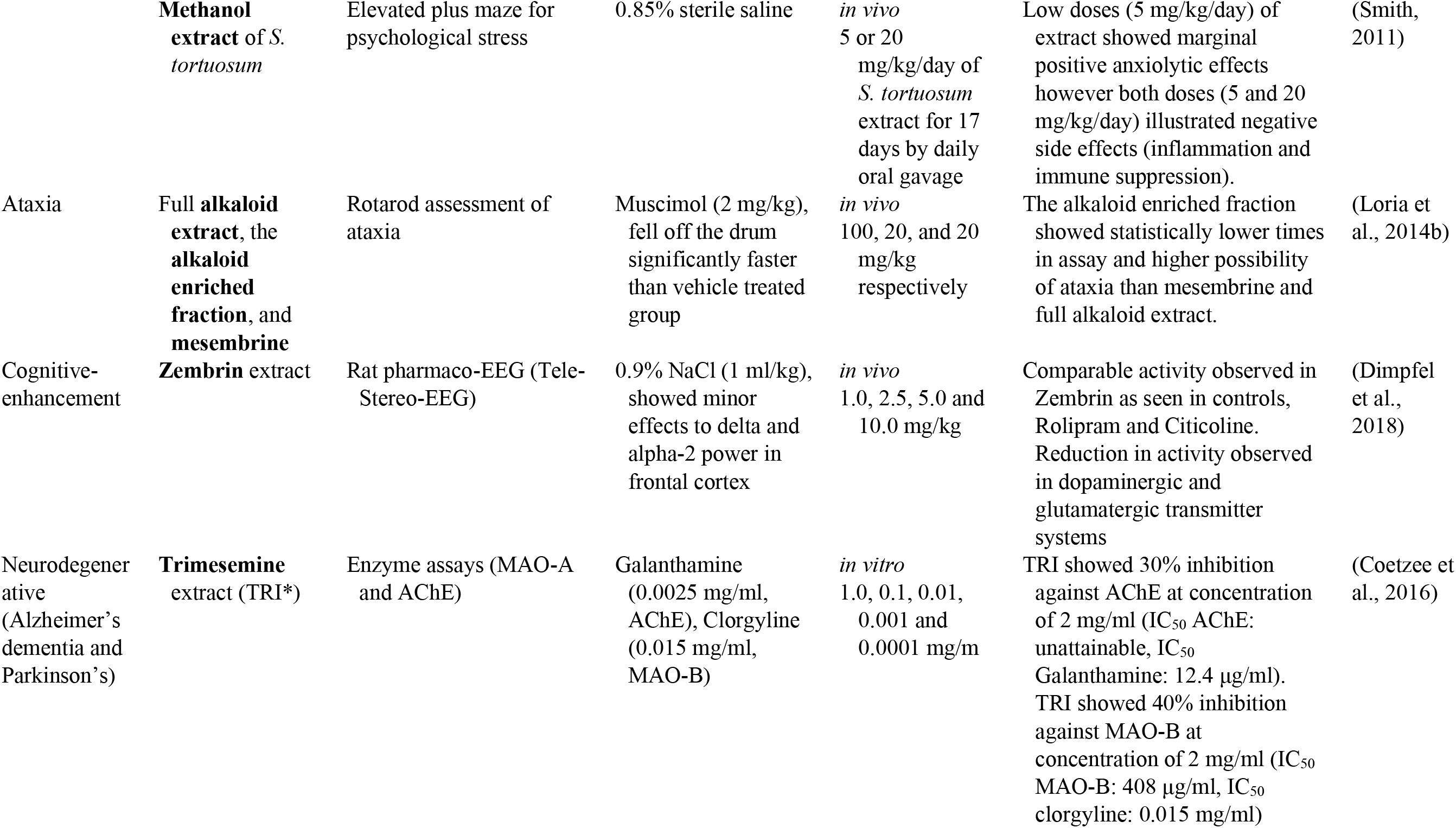
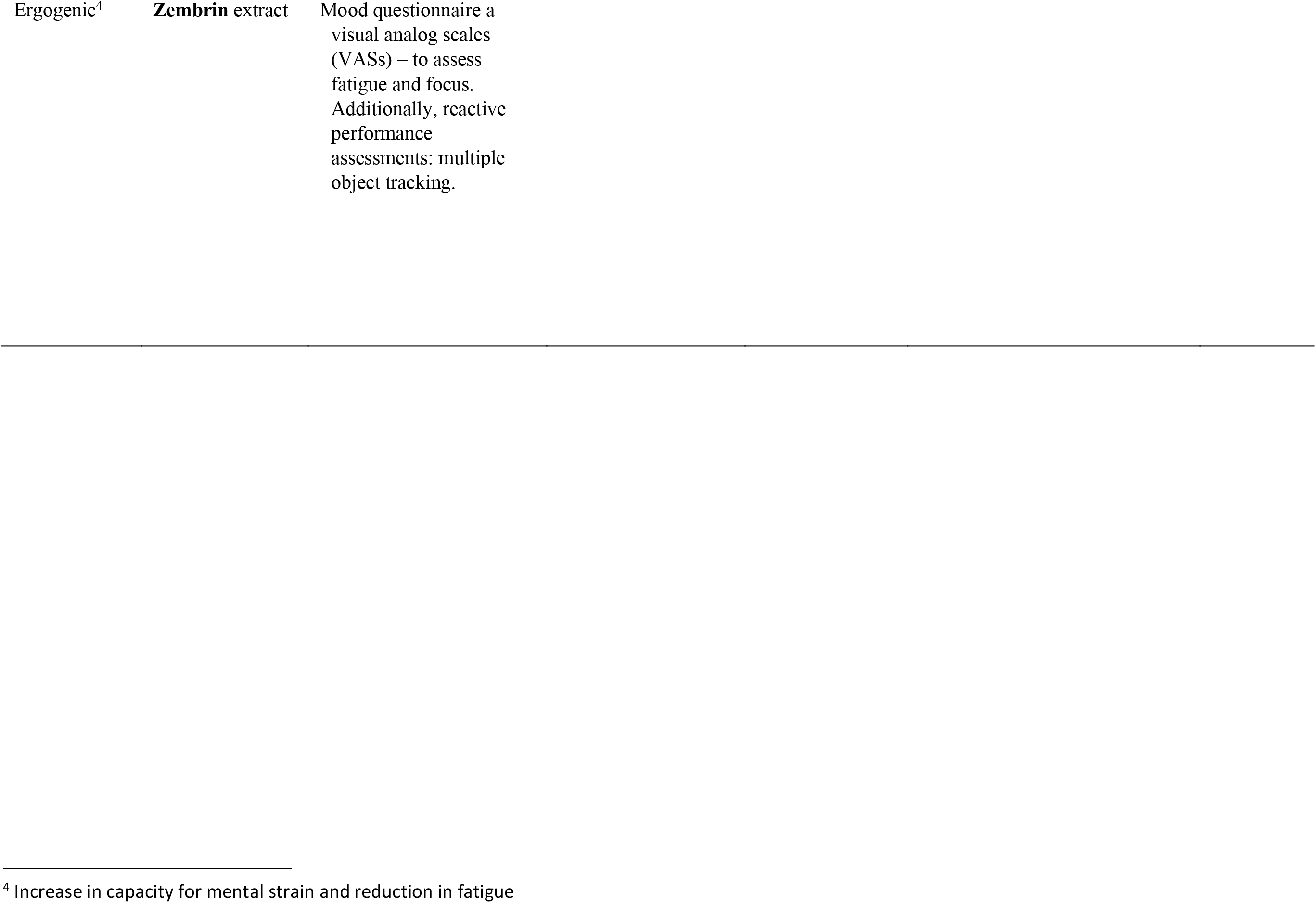
The CNS-related activity of Sceletium tortuosum extracts and compounds (*Note that TRI is an extract of a S. tortuosum and S. expansum hybrid)

Bennett and Smith, (2018) also report that the high-mesembrine *Sceletium* extract, Trimesemine™ could hold a potential therapeutic activity in cytokine induced depression as a preventative supplement. They propose that the extract modulates the basal inflammatory cytokine profile whilst maintaining that, there is no change in the acute response to pathogenic challenge (Bennett and Smith, 2018). Furthermore, these findings are an indication that this extract may prove to be of direct benefit to the attenuation of cytokine-induced depression, as well as in systemic low-grade inflammation in immune cells (Bennett and Smith, 2018). This particular study did not test individual alkaloids, and such could not pinpoint the plant constituent responsible for the observed activity. In the future, extraction and isolation of alkaloids may prove beneficial if the intention is to correlate bioactivity to specific alkaloid constituents so that our overall understanding of which phytochemicals are responsible for the observed activity can be clarified.

Receptor screening of Zembrin® (a standardised extract of *Sceletium tortuosum*) was conducted against 77 radioligand binding assays (0.75 mg/mL and a panel of phosphodiesterases) to compile a comprehensive list of potential CNS and other pharmacological targets (Harvey et al., 2011). The extract showed binding at the serotonin (5-HT) transporter, δ2- and μ-opioid receptors, the cholecystokinin-1 receptor, > 80% inhibition at GABA receptors (non-selective) and PDEs 3 and 4 (Harvey et al., 2011). Some of the therapeutic applications of these targets are emesis, obesity, anxiety and migraine linked to the serotonin (5-HT) transporter (Pithadia and Jain, 2009). The δ2- and μ-opioid receptors are involved in maintaining epileptic seizure, emotional responses, immune function, obesity, cell proliferation, respiratory and cardiovascular control as well as several neurodegenerative disorders (Feng et al., 2012). The cholecystokinin-1 receptor is involved in gastrointestinal and metabolic diseases (Berna and Jensen, 2007). The GABA receptors are involved in pathologies ranging from epilepsy, schizophrenia, anxiety disorders and premenstrual dysphoric disorder (Wong et al., 2003). The PDE3 receptor is responsible for platelet activation/aggregation (Beca et al., 2011), and vascular smooth muscle proliferation (Beca et al., 2011; Begum et al., 2011), while the PDE4 receptor is responsible for inflammatory conditions including asthma, chronic obstructive pulmonary disease (COPD), psoriasis, atopic dermatitis (AD), inflammatory bowel diseases (IBD), rheumatic arthritis (RA), lupus, and neuroinflammation (Li et al., 2018). This report only presented findings on the affinity of Zembrin® to different receptors. Further studies would need to be investigated for activity against these specific pathologies. Plant extracts have numerous metabolites that work in synergy to affect their biological influence and it is thus possible that multiple metabolites are potentiating the mood-elevated activity aside from mesembrine alone, as suggested by Lubbe et al. (2010). They emphasised that irrespective of fermentation, the crude extracts of *S. tortuosum* showed greater inhibition of acetylcholinesterase than mesembrine alone. This could be indicative of the possibility that other phytochemicals may be involved in potentiating mood-elevating activity (Lubbe et al., 2010).

The Harvey et al. (2011) study supports ethnobotanical use as a mood-elecator by the observed serotonin transport activity in response to Zembrin®. The serotonin receptor influences a myriad of biological and neurological processes such as anxiety, appetite, aggression and depression (Mück-Šeler and Pivac, 2011; Zhang and Stackman, 2015). Evidence of anxiolytic effects of *Sceletium* in humans (Gericke and Viljoen, 2008) has partially been supported in a study using a rat model of restraint induced stress (Smith, 2011). The binding of compounds to various sites on the 5-HT transporter (SERT) is considered evidence of potential serotonin reuptake inhibition, a common target of antidepressant drugs. A selection of alkaloids, mesembrine, mesembrenone, and mesembrenol, from *Sceletium tortuosum,* were tested for their affinity for SERT with *K*i*’s* of 1.4, 27, and 63 nM, respectively (Harvey et al., 2011). These values were significantly higher than other alkaloids, such as buphanidrine or distichamine, isolated from Amaryllidaceae, with reported *K*i’s of 312 and 868 μM, respectively (Neergaard et al., 2009). These compounds have been found to already possess well-established anti-depressant activity, found in *Boophone disticha* (L.f.) Herb (Amaryllidaceae) (Neergaard et al., 2009). There is some evidence that argues against *Sceletium* purely acting as a selective serotonin re-uptake inhibitor (SSRI), as repeated administration of SSRIs has been linked to hyposensitivity to SSRIs as a result of an up-regulation in PDE4 (Ye et al., 2000). However, it has been demonstrated that PDE4 activity decreased after *Sceletium* administration (Harvey et al., 2011).

Harvey et al. (2011) study was the only study testing *S. tortuosum* extract on a number of receptors. More studies should be done investigating other species of *Sceletium* and isolated compounds on an array of receptors suspected to control anxiety. Although the Harvey et al. (2011) study looked at the pharmacokinetics of individual compounds against each receptor, it may be valuable to test extracts with varying concentrations of compounds to test samples more aligned with ethnobotanical use.

*In vivo* testing of *Sceletium* alkaloids has been performed using rat models designed for mental disorders such as neurodegeneration, like Alzheimer’s Disease (AD), epilepsy and depression (Loria et al., 2014a) (Fig. 5c). Loria et al. (2014a) found that mesembrine from *S. tortuosum* had analgesic and antidepressant activity. *Sceletium* species have exhibited potential therapeutic activity *in vivo* using rodent models for AD, anxiety and depression (Gericke and Viljoen, 2008; Krstenansky, 2017). A summary of the CNS-related activity, together with the recent anti-inflammatory activity of *Sceletium* is presented in Table 3. With the PDE4 activity of *Sceletium* extracts noted by Harvey et al. (2011) and new *in vivo* on the receptor itself, there is evidence suggesting that inhibitors from *Sceletium* can aid to reverse depression, improve cognitive ability and reduce anxious states. Ataxia and cognitive-enhancement potential of *Sceletium* extracts and Zembrin medications are also reported in Table 3. Several tests have been conducted on animals at a significantly larger dose than has been reported as the recommended dose via tablets or capsules (1-2 mg/kg per day) (Hirabayashi et al., 2002). Furthermore, the study also found that the antidepressant activity of Zembrin was unfortunately associated with ataxia^5^ (Hirabayashi et al., 2002).

The anxiolytic activity of *Sceletium* may be attributed to other mechanisms besides serotonin-reuptake inhibition such as monoamine release (Coetzee et al., 2016). Hirabayashi et al. (2002)., found that a mesembrine extract primarily had anti-depressant and anxiolytic activity associated with the activity of the monoamine oxidase system. They also reported on mild toxicity and the potential for the high mesembrine *Sceletium* extract (Trimesemine^TM^) for other neurological disorders such as Alzheimer’s disease and attention deficit disorders (Coetzee et al., 2016)

*In vivo* testing of *Sceletium* alkaloids has been performed using rat models designed for mental disorders such as neurodegeneration, like Alzheimer’s Disease (AD), epilepsy and depression (Loria et al., 2014a) (Fig. 5c). Loria et al. (2014a) found that mesembrine from *S. tortuosum* had analgesic and antidepressant activity. *Sceletium* species have exhibited potential therapeutic activity *in vivo* using rodent models for AD, anxiety and depression (Gericke and Viljoen, 2008; Krstenansky, 2017). A summary of the CNS-related activity, together with the recent anti-inflammatory activity of *Sceletium* is presented in Table 3. With the PDE4 activity of *Sceletium* extracts noted by Harvey et al. (2011) and new *in vivo* on the receptor itself, there is evidence suggesting that inhibitors from *Sceletium* can aid to reverse depression, improve cognitive ability and reduce anxious states. Ataxia and cognitive-enhancement potential of *Sceletium* extracts and Zembrin medications are also reported in Table 3. Several tests have been conducted on animals at a significantly larger dose than has been reported as the recommended dose via tablets or capsules (1-2 mg/kg per day) (Hirabayashi et al., 2002). Furthermore, the study also found that the antidepressant activity of Zembrin was unfortunately associated with ataxia (Hirabayashi et al., 2002). A new structure-function relationship for *Sceletium* alkaloids was suggested by Timoneda et al. (2019), tests performed on rats using Zembrin® found new evidence of electric excitability of the rat hippocampus supporting this new relationship.

**Table 2:**
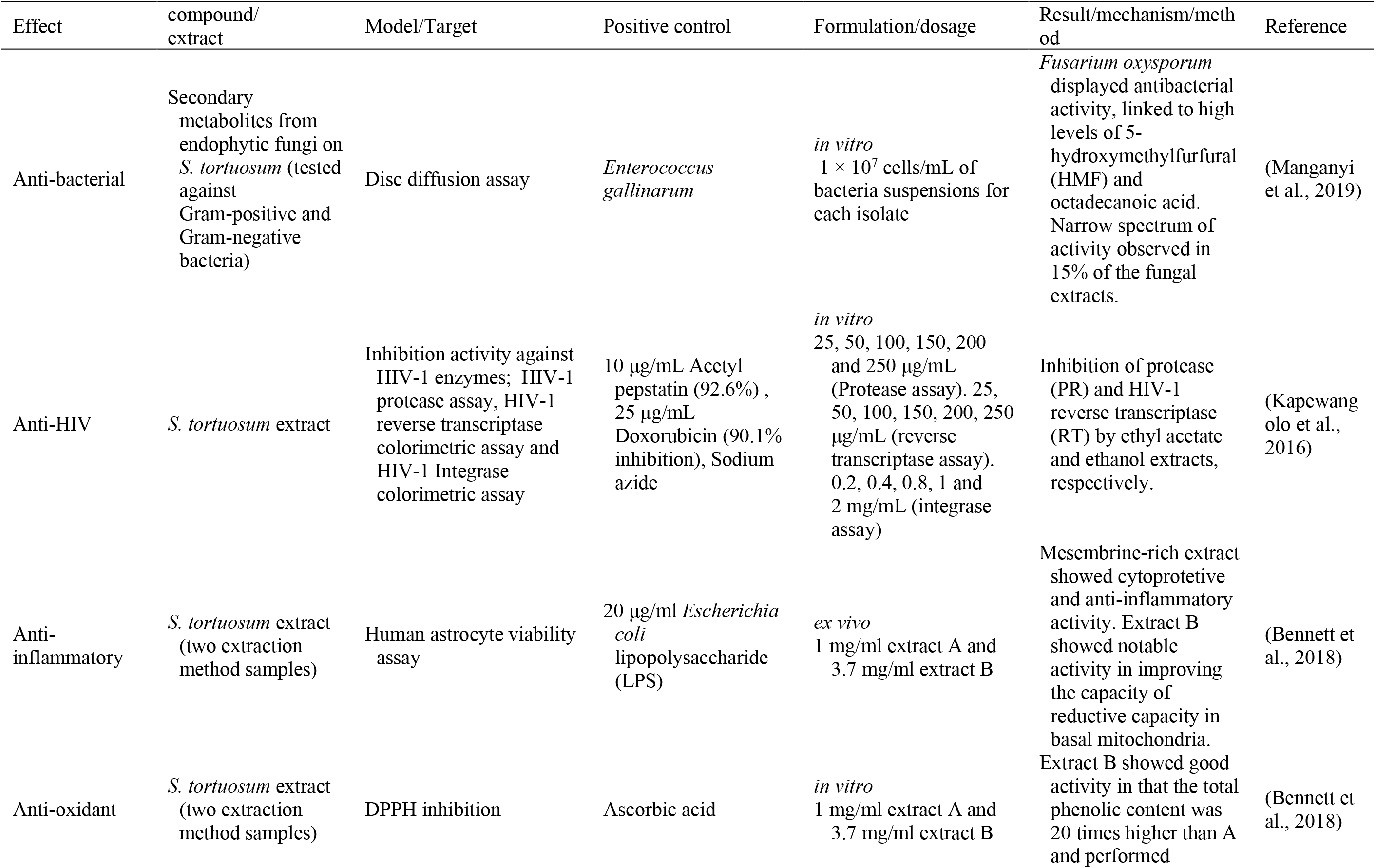
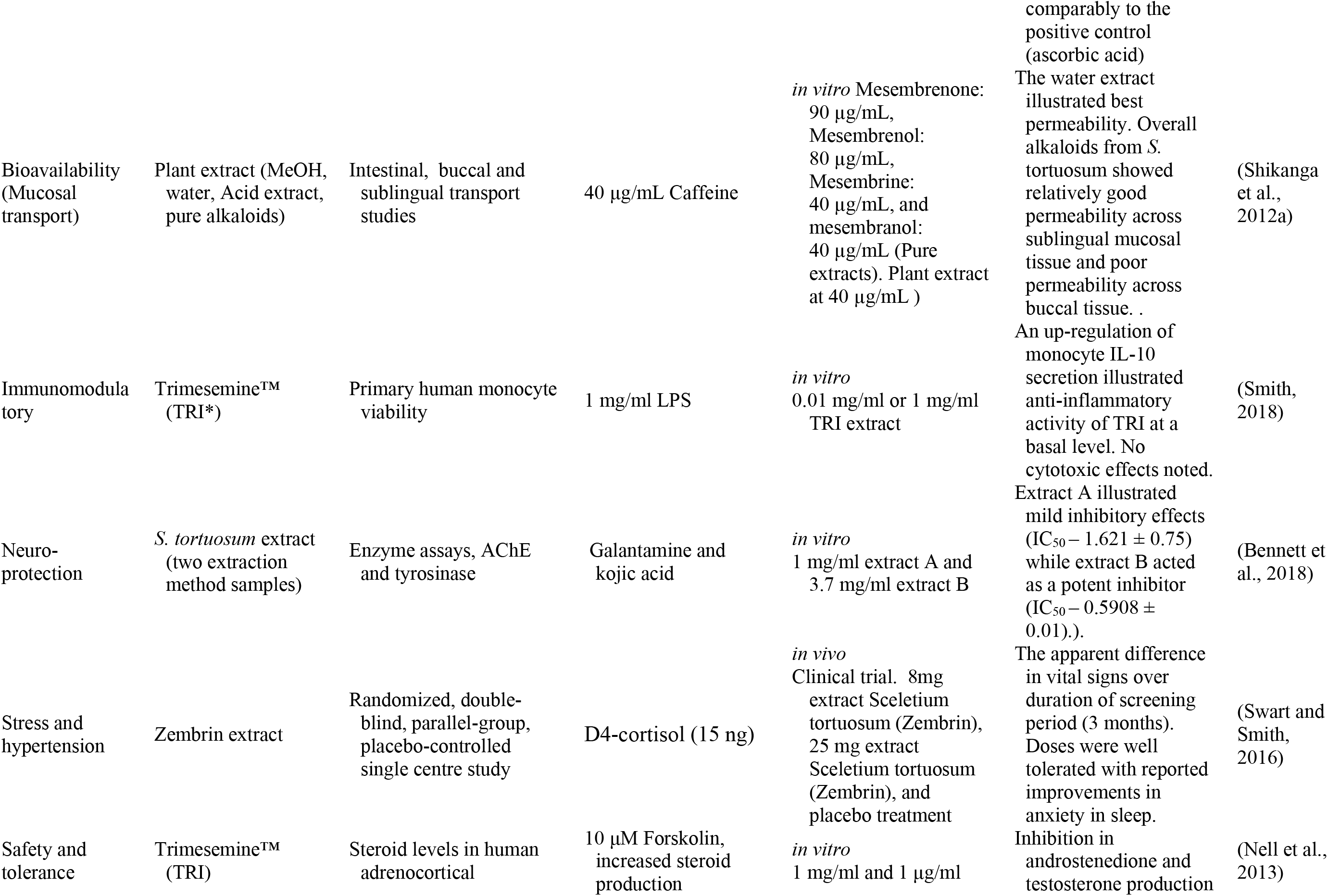
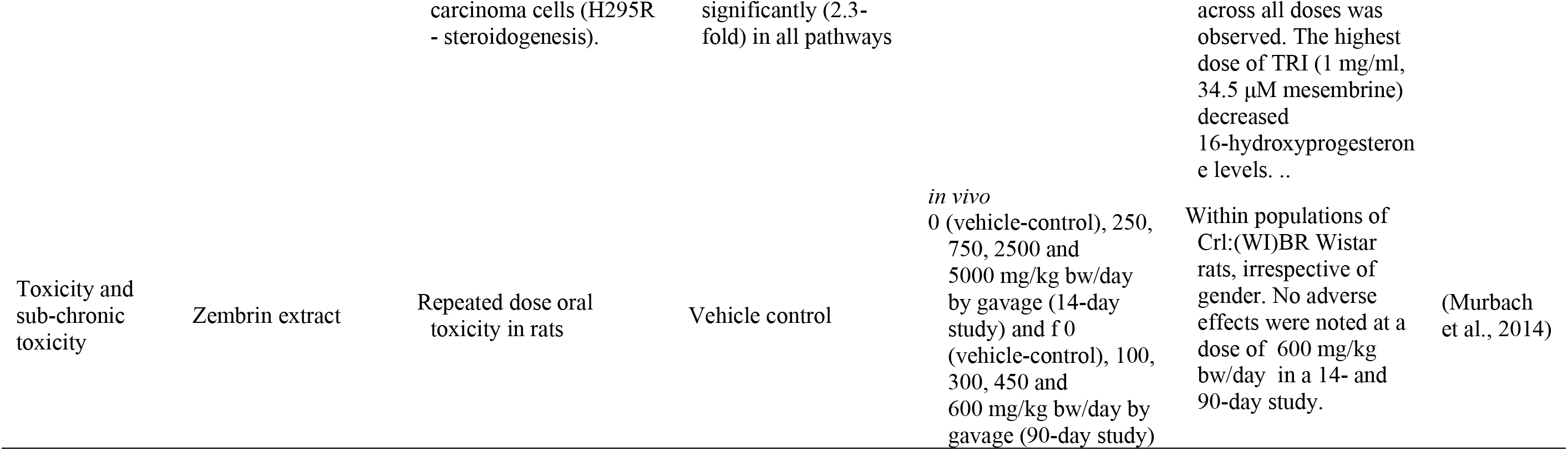
Other notable biological activity of Sceletium tortuosum extracts and compounds (*Note that TRI is an extract of a S. tortuosum and S. expansum hybrid

Mesembrenone has shown anti-tumour activity with cytotoxicity being tested against a murine non-tumoral fibroblast cell line and a human tumoral cell line (Molt4) (Weniger et al., 1995). Weniger et al. (1995) tested 25 alkaloids from Amaryllidaceae with only mesembrenone showing some specificity for Molt4 cells. Recently an investigation by Bennett and Smith, (2018), on the anti-inflammatory potential of *Sceletium,* reported that extracts high in mesembrine and Δ^7^mesembrenone exerted anti-inflammatory and antioxidant activity *in vitro* respectively. The mesembrine-rich extract, which was less refined as compared to the Δ^7^mesembrenone extract, exhibited broad dose range efficacy and may serve as a promising therapeutic in the setting for chronic diseases (Bennett et al., 2018). The Δ^7^mesembrenone rich extract exhibited good antioxidant activity which was noted as safe at low doses of administration (Bennett et al., 2018). Due to the aetiology of both diabetes and obesity being linked to inflammation and excess glucocorticoid production (Bennett et al., 2018), these findings may hold value in chronic lifestyle disease management such as diabetes, type-1 and -2. For such data to be translated into a pharmaceutical drug discovery chain, endocrine-immune (IL-6 and MCP-1) interactions need to be investigated.

The potential therapeutic activity of mesembrine alkaloids towards inflammatory diseases ranging from asthma, chronic obstructive pulmonary disease, psoriasis, and, treating depression has been investigated (Houslay et al., 2005). The anti-inflammatory activity is suspected to be due to the activity of mesembrine-HCl acting as an inhibitor of phosphodiesterase-4 (PDE4), with observed activity at an IC_50_ of 29 µM (Napoletano et al., 2001). Through the selective inhibition of the PDE4 family of enzymes is predicted to generate great functional effects, as evidenced by PDE4 enzymes being a major therapeutic target for inflammatory diseases (Gericke and Viljoen, 2008). Phosphodiesterase-4 (PDE4) is the major class of enzymes that mediates the hydrolysis of cyclic adenosine monophosphate (cAMP). PDE4 has also been identified to play a role in the inflammatory system (Banner and Trevethick, 2004; Dastidar et al., 2007; Li et al., 2018). Harvey et al. (2011) found that the alkaloids mesembrenol, mesembrine and mesembrenone inhibited PDE4B with IC_50_ values of 16, 7.8, and 0.47 μM, respectively. The positive control, rolipram had a IC_50_ for PDE4B of 0.13 μM (MacKenzie and Houslay, 2000). Another enquiry illustrated the activity of Zembrin® *in vivo* to correlate this mechanism of action observed *in vitro* in freely moving rats interpreted as an electopharmacogram^6^ (Dimpfel et al., 2016).

More recent studies include the work of Reay et al. (2020), where the anxiolytic properties of Zembrin® were assessed in a double-blind, placebo-controlled behavioural study with healthy human volunteers. A dose of 25 mg was administered and stress was assessed. The volunteers consisted of younger human adults, who were subjected to two assessments of stress. Namely, a multitasking framework and a simulated public speaking framework. The results of this study indicated that 25 mg of Zembrin® tentatively illustrates the anxiolytic properties of *S. tortuosum*. However, the study fails to replicate previously reported enhancement of cognitive function. They note this absence of cognitive enhancement to be the first evidence of Zembrin® having no impact on nonexecutive memory processing in healthy participants.

Maphanga et al. (2020), assessed the anxiolytic activity in a zebrafish behavioural assay for a number of medicinal plants one of which was *S. tortuosum*. The assay consisted of using 5-day post-fertilized zebrafish larvae and monitoring their movements (spontaneous locomotor activity) in response to light-dark transitions used to induce an anxious state. Whole plant parts were used and a distilled water extract was made. The only extract to show a significant (p < 0.001) reduction in anxiety (reverse-thigmotaxis assessment) was *S. tortuosum*, as compared to the control group. The extract was then subjected to more sophisticated anxiolytic assessments where it was observed that the concentrations of 12.5 and 25 mg/L showed significant levels in reducing the anxiolytic state. Additionally, no toxic effects were observed on the zebrafish in the assay. The model proves to be an appropriate and repeatable assay for assessing the anxiolytic activity of *Sceletium* extracts. Supporting this work is the rising application of the zebrafish animal model as it is claimed to be adequately comparable to humans, sharing approximately 70-80% genetic homology with humans (Barbazuk et al., 2000; Goldsmith, 2004).

The ergogenic effect of *S. tortuosum* has been investigated in a clinical trial with human participants investigating the potential of *S. tortuosum* as a supplement to reduce fatigue and improve focus. The study was conducted on men and women over 8 days with supplementation of the *S. tortuosum* commercially available extract, Zembrin (Hoffman et al., 2020). In the subject population studied, no benefits in mood were observed. However, there were significant improvements noted in complex reactive performance tasks that include the stress of cognitive load. It should be noted that other forms of assessments linked to measuring cognitive and mood information may yield different results. Furthermore, no pharmacokinetics and absorption data are presented in the study.

Zebrafish assays to assess the anxiolytic activity of *S. tortuosum* alkaloids were once again studied by Maphanga et al. (2022). The study now assessed isolated alkaloids from the extract described in Shikanga et al. (2011), at concentration ranges of 10. 15, 30 and 50 μM with the greatest activity across alkaloids observed at 50 μM. Toxicity measured as MTCs (Maximum Tolerated Concentration) where locomotor activity was impaired in the zebrafish was observed at concentrations between 75 and 150 μM and the MTC was identified at 50 μM (Maphanga et al., 2022).

Gericke et al. (2022), assessed the acute antidepressant-like effects of Zembrin in a Flinders Resistant line (FSL), a rodent model for depression. The study reported that Zembrin at doses of 25 and 50 mg/kg were effective antidepressants in the forced swim test (FST) and performed better than the control (Escitalopram). This was the first study to date that compared Zembrin to an SSRI in a rodent model of this kind, supporting the therapeutic use of *S. tortuosum* for mood disorders.

Bioavailability studies on *Sceletium* and its alkaloids are greatly lacking in research. Shikanga et al., (2012a), presented findings on the permeability of mesembrine across the buccal, intestinal and sublingual mucosal membranes. In that study, mesembrine had a higher permeability across intestinal tissue than the positive control caffeine but the permeability was lower in the buccal and mucosal sublingual membranes. Manda et al. (2017), showed that the oral bioavailability of mesembrine and mesembrenone in mouse plasma (using UHPLC-QToF-MS) was poor and below the detection limit. Bioavailability information regarding other alkaloids and chemotypes from *Sceletium* are still not documented in terms of data on the permeability of these alkaloids across buccal, intestinal and sublingual mucosal tissues. It is thus imperative that more attention should be placed on such to provide new evidence linked to bioavailability in order to support or refute ethnobotanical claims. It may also be of interest to monitor cultivated and commercially available samples for pesticide residues and toxic alkaloids in other plants that may have mistakenly been gathered during the harvesting of wild populations of *S. tortuosum* as this species is often found under the canopy of other small shrubs in the wild and in close association with a diverse range of other species. At this present time, there is no information in this respect and the monitoring of plant or chemical contaminants is thus urgently needed.

### Biological associations

Recently, a new avenue of investigation was based on investigating the association of endophytic fungal communities on *S. tortuosum* (Manganyi et al., 2018) (Fig 5C and D). *Fusarium, Aspergillus* and *Penicillium* were among the fungal endophytes found in the plant. In total there were 60 endophytic fungal species successfully isolated and identified, belonging to 16 genera. The antibacterial activity of this endophytic fungi was also investigated, where it was found that some fungal isolates could provide sources of novel antimicrobial agents against anti-biotic resistant strains (Manganyi et al., 2019). This is also the first investigation to report on secondary metabolites from endophytic fungi, *F. oxysporum* (GG 008, accession no. KJ774041.1) isolated from *S. tortuosum* (Manganyi et al., 2019).

### Propagation techniques

Faber et al. (2020), reported on the influence of soilless growth medium (pure silica sand, 50% silica sand with 50% coco-peat, 50% silica sand with 50% vermiculite, and 50% silica sand with 50% perlite) and fertigation regimes (nutrient solution administered in intervals from 1-5 weeks) on shoot and root growth as well as how these factors influenced alkaloid levels (D7-mesembrenone and mesembrine). The study concluded that shoots had higher concentrations of mesembrine while roots had higher concentrations of mesembrenone and d7-mesembrenone. Shoots overall had higher alkaloid concentrations than that of shoots. Fertigation results vary in plant parts and across alkaloids, thus further work is needed to this end. The major observation is that the influx of secondary metabolites in *S. tortuosum* seems to respond to biotic and abiotic factors (Bourgaud et al., 2001; Ashraf et al., 2018).

To date, there have only been three studies investigating the micropropagation of the medicinally important, *S. tortuosum* (Sreekissoon et al., 2021b, 2021a; Makunga et al., 2022). The illegal harvesting and exploitation of *S. tortuosum*, due to the demand as a recreational drug linked to mood-elevating properties of the mesembrine alkaloids is a driving factor for the dire need for the development of micropropagation techniques. Development of micropropagation techniques with *Sceletium* could offer a direct and standardised source of mesembrine alkaloids. Sreekissoon et al., (2021b) investigated whether *in vitro* regeneration of micropropagules with auxins could be acclimatised *ex vitro*. Sreekissoon et al. (2021a), assessed the effects of smoke water on the germination, seedling vigour and growth of *S. tortuosum*, the major findings were that seed vigour was highest at smoke water concentrations of 1:1000. A limitation of these studies was that key biomarker compounds were not monitored in the microplant regenerates. The Makunga et al. (2022), study investigated morphotypes of *S. tortuosum* and the levels of mesembrine and its derivatives. For the first time, the alkaloids, Δ4-mesembrenone, mesembrenol, mesembrine, and mesembranol were reported in the literature (Makunga et al., 2022). The utilisation of micropropagation using the dehydrating and rehydration technique outlined in this report, resulted in *in vitro* metabolite accumulation comparable to wild-type material collected (Zhao et al., 2018a).

### Legislation, toxicology and safety of *Sceletium* alkaloids

Toxicological assessments on *Sceletium* are limited with the first formal *in vivo* toxicological assay being performed by Murbach et al. (2014) on the mesembrine-rich extract Zembrin^®^. They found that Zembrin^®^, in male and female Crl:(WI)BR Wistar rats, showed no mortality or treatment-related adverse effects spanning 14 or 90 days with doses of 600 and 5000 mg/kg bw/day, respectively (Murbach et al., 2014). A greater effort in understanding cytotoxic effects of *Sceletium*-derived extracts and their potential drug-herb interactions is also urgently needed.

Clinical administration of *Sceletium* tortuosum has been carried out by Gericke (2001). The clinical case study reported on three individuals who have been prescribed *S. otrtuosum* in tablet form. The first individual was initially suffering from weight loss, appetite loss, anxiety, insomnia and severe depression that had been ongoing for 4 months. A dose of 50 mg/day was prescribed and reported an initial increase in anxiety for about a week. After which, symptoms of anxiety ceased to show and a general improvement in mood was reported. The tablets were stopped after 4 months, with no apparent signs of withdrawal symptoms.

The second individual was a patient with a diagnosed personality disorder (dysthymia). The patient experienced feelings of anxiety, depression and hypersomnia. The patient was prescribed 50 mg tablets for 10 days, after which upon request the dose was increased to 2 tablets (100 mg). The patient described an overall decrease in anxiety and was more able to cope with stress in her life.

The third individual presented with major depressive disorder. The symptoms presented were over-eating, anxiety, depression, a general lack of motivation and suicidal thoughts. The patient was prescribed 2 tablets (100 mg) per day. Symptoms of anxiety and depression were absent within the first day. The course was maintained for 6 weeks after which, no signs of withdrawal symptoms were present.

The plant is marketed as a food supplement and not as a scheduled drug. The United Nations Office on Drugs and Crime (UNODC) flagged *Sceletium* as a plant of concern when reporting on substances of concern in 2013 as part of a report on the obstacles in the identification and regulation of new psychoactive substances (UNODC, 2013). Due to plant material being crushed upon commercial sale, it is difficult to authenticate plant material. The nature of *Sceletium* being classified as herbal or dietary supplements often exempts it from mandatory testing by the US Food and Drug Administration (FDA). Thus, without adequate quality control and authenticating systems in place, herbal product producers can adulterate samples. This has been observed with commercial *Sceletium* products in Germany, where plant material was laced with the stimulant ephedrine (which has been banned in herbal products and supplements) (Lesiak et al., 2016).

Dietary supplementation is popular for those that partake in sports recreationally or as professional athletes and sports performance-enhancing natural products are thus highly sought after. Legislation around the utilisation of *S. tortuosum* extracts for elite athletes in competitive sports as a dietary supplement hangs in the balance as some regulatory bodies have denoted its status under the categories of ‘unauthorized novel food’ and it is included also in the European Food Safety Authority Compendium of Botanicals as concerning for human consumption (Jędrejko et al., 2021). Due to its effects on brain function and cognition, it has not necessarily been approved for routine use by the World Anti-Doping Agency (WADA) which regulates permissible dietary supplements for athletes.

### Conclusions and future prospectives

A comprehensive review of the literature together with bibliometric meta-data analysis has identified the gaps and achievements in research on this important southern African medicinal genus, presented in this review. The bibliometric analysis showed that South Africa has established a strong network with researchers working on *Sceletium* and its medicinal value but there is an apparent lack of synergy and coordination between research groups located in the native land of this genus. The reason for this is not obvious but could be linked to historical networks and collaborations being preferred and strong competition for limited funding between research groups. A higher degree of collaboration is thus foreseen to encourage greater progress in transforming latent botanical assets into consumer products.

A portion of the current review has highlighted the development of quality control tools for commercial and wild-harvested *Sceletium* species, with the focus being on *S. tortuosum* due to its rising commercialisation status in global markets. With the popularity of *Sceletium* growing as a recreational natural product, a more diverse range of products emanating from a growing number of manufacturers is an imminent probability. However, the expansion of the industry may bring about an increased frequency of herb-drug adulterations as seen with many other natural products that are popular in commercial settings (Campbell et al., 2013; Seethapathy et al., 2015; Booker et al., 2016). There is thus a critical urgency that is required in the research and development of analytical techniques and protocols that are rapid, robust and reproducible which may be highly efficacious in their detection capacity for adulterants that may occur in products even at minute scales. The inconsistent chemical profiles and various wild chemotypes of *S. tortuosum* and its sister species in the wild (Zhao et al., 2018a) further necessitate the application of techniques for quality control analysis that have high resolving power. Such techniques need to be able to identify adulterants that would otherwise not be identified due to their similarity in polarity with some of the biomarker compounds. From a practical perspective, these instruments are not always common in laboratories and they are expensive, requiring users to have specialized training and highly-sophisticated scientific expertise.

The choice of chemo-elite types that can become easily domesticated may assist the production of quality-assured natural products, generating an industry that will gain consumer trust. This is viewed as being of high importance when considering that some *S. tortuosum* wild types may produce mesembrine alkaloids at exceedingly low concentrations and other *Sceletium spp.* show a complete lack of the key biomarker compounds that are routinely examined by the phytopharmaceutical and nutraceutical industries (Patnala and Kanfer, 2013). Because the different species are similar to each other, this makes them highly vulnerable to misidentification and incorrect identification and unregulated collection of these plants may set in motion their overharvesting thereby creating serious conservation concerns of the genus. The number of studies looking into genetic approaches for quality control tools is remarkably absent in *Sceletium* research. DNA fingerprinting and biomarker identification (Klak et al., 2007; Shikanga et al., 2013b; Zhao et al., 2018), single nucleotide polymorphisms (SNPs) and microsatellite loci (nuclear short sequence repeats, SSR) have been commonly used for genetic-based quality control (Laurie et al., 2010). The absence of quality pure chemical standards has greatly hindered the absolute quantification of the mesembrine alkaloids in many analytical studies that have been performed on *Sceletium*. More effort is thus required to fill this gap as the lack of reference compounds makes the identification and profiling of both minor and major alkaloids synthesised by *S. tortuosum* and its relatives more challenging.

Historical ethnobotanical records allude to a practice of fermentation of S. tortuosum when it is used by local indigenous people but scientific evidence of the fermentation on phytochemicals of the plants remains in contention as current fermentation studies do not correlate with each other. Studies investigating the manipulation of biosynthetic pathways in combination with fermentation studies may thus prove valuable in deepening the general understanding of the effects of the fermentation treatment(s) and its biological effects in animal systems. Despite this, the pharmacological tests, whether they be in *in vitro* and/or *in vivo* experiments, have increasingly supported the traditional use of *S. tortuosum* as a mood-elevator and anxiolytic agent. However, its anti-inflammatory and immunomodulatory effects have been insufficiently investigated up until recently. At this point, there has been some evidence that highlights the beneficial physiological effects of *S. tortuosum* extracts as a plant medicine which extend beyond its psychoactive effects, with potential therapeutic activity targeted at diabetes and obesity (Bennett and Smith, 2018). Interest in developing *Sceletium* into an additive for foods, beverages and supplements aiding in depressive and anxiolytic disorders has been happening for over a decade (Gericke and Viljoen, 2008), with some products finding the market. If this is to be fully realised, a fundamental field of investigation that will need to be looked into is the standardised cultivation of the plants for their phytochemicals. This can be achieved through the manipulation of secondary metabolites using physiological stress such as light, pH and nutrient stress in *Sceletium* species, which is currently a void in the research scope.

To enable such studies, a reference genome is also urgently needed for *S. tortuosum* as currently the genetic resources that may assist with understanding the genetic and biochemical controls that are involved in the biosynthetic pathways of mesembrine alkaloids are unavailable. Such resources would thus provide additional research efforts into the control of metabolic flux linked to mesembrine biosynthetic pathways and identify regulatory promoters that influence the synthesis of the unique alkaloids of *Sceletium*. Systems biology studies using a multi-omics approach may further assist with the full characterization of pathway interactions that may lead to a better understanding of the metabolic networks that control alkaloid biosynthesis routes of *Sceletium tortuosum* and related sister species.

Ultimately, *Sceletium* and its alkaloids hold great potential in future endeavours and might provide novel insights into the synthesis pathways of *Sceletium*-specific alkaloids and their genetic regulatory controls whilst studies on inflammation activity and phytomedicinal applications of the plant rise in industry.

## Ethics declarations

### Availability of data and material

All data generated for this review has been included in the manuscript.

### Authors’ contributions

KR conducted the bibliometric analyses and wrote the first draft of this manuscript. GIS and NPM conceptualized the study and contributed by editing the draft versions of this paper. All authors read and approved the final version of this review article.

## Footnotes

5 Impaired balance or coordination can be due to damage to brain, nerves or muscles.

6 An analysis recording the field potentials in the frontal cortex, striatum, hippocampus and reticular formation while administered drugs.

